# Mitochondrial effects on fertility and longevity in *Tigriopus californicus* contradict predictions of the mother’s curse hypothesis

**DOI:** 10.1101/2022.04.20.488069

**Authors:** Eric T. Watson, Ben A. Flanagan, Jane A. Pascar, Suzanne Edmands

## Abstract

Strict maternal inheritance of mitochondria favors the evolutionary accumulation of sex-biased fitness effects, as mitochondrial evolution occurs exclusively in female lineages. The “mother’s curse” hypothesis proposes that male-harming mutations should accumulate in mitochondrial genomes when they have neutral or beneficial effects on female fitness. Rigorous empirical tests have largely focused on *Drosophila*, where support for the predictions of mother’s curse has been mixed. We investigated the impact of mother’s curse mutations in *Tigriopus californicus*, a minute crustacean. Using nonrecombinant backcrosses, we introgressed four divergent mitochondrial haplotypes into two nuclear backgrounds and recorded measures of fertility and longevity. We found that the phenotypic effects of mitochondrial mutations were context-dependent, being influenced by the nuclear-background in which they were expressed, as well as the sex of the individual and rearing temperature. Mitochondrial haplotype effects were greater for fertility than longevity, and temperature effects were greater for longevity. However, in opposition to mother’s curse expectations, females had higher mitochondrial genetic variance than males for fertility and longevity, little evidence of sexual antagonism favoring females was found, and the impacts of mitonuclear mismatch harmed females but not males. Together, this indicates that selection on mitochondrial variation has not resulted in the accumulation of male mutation load in *Tigriopus californicus*.

## Introduction

In eukaryotes, life is powered by mitochondria - cytoplasmic organelles capable of producing energy in the form of ATP. Originating from an ancient symbiosis over 2 billion years ago, mitochondria provided the energy required for the evolution and diversification of multicellular organisms[1–3]. Over evolutionary history, mitochondrial genomes have retained some aspects of their ancestral genetic codes and a few dozen genes while dramatically reducing their genome size through deletion and gene transfer to the nucleus. Beyond the generation of ATP by oxidative phosphorylation (OXPHOS), mitochondria perform additional cellular functions, including initiation of apoptosis, amino acid and steroid biosynthesis, β-oxidation of fatty acids, cellular signaling, and nuclear epigenetic regulation [4–8]. Nevertheless, out of 1,500 or more proteins found in the mitochondrion, only the 13 membrane-bound electron transport chain complexes are encoded by the mitochondrial genome, with the rest being encoded by nuclear DNA[9,10]. The coevolution of mitochondrial and nuclear genomes is thus an ancient yet ongoing process, involving the evolutionary fine-tuning of various molecular interactions (i.e. protein-protein, protein-DNA, and protein-RNA) required for the accurate replication and transcription of mtDNA, translation of mtDNA-encoded proteins, and efficient cellular respiration. Mutation rates are generally higher in mtDNA than in nDNA[11,12] leading to a continuous introduction of new mitochondrial alleles. With asexual transmission and lack of recombination in mitochondria, novel alleles are likely to persist even when slightly deleterious due to the potential masking effect of linkage disequilibrium on mitochondrial genetic variation and reduced efficacy of natural selection[13], although debate over the efficacy of selection on mtDNA continues to produce compelling counterarguments (see [14–18]). Nevertheless, mtDNA likely experiences recurrent adaptive evolution via selective sweeps acting on traits such as respiratory efficiency and mitonuclear coadaptation[19–23]. In natural populations, mitochondrial fitness variation is likely to be present at substantial levels and may make a meaningful contribution to adaptation and diversification[24]. The distribution of fitness effects among mtDNA mutations is also likely to differ between the sexes, owing to the maternal inheritance of mitochondria[25–27]. Strict maternal inheritance favors the evolutionary accumulation of sex-biased fitness effects, as it is expected to preclude any opportunity for evolution in males, forming a sex-specific selective sieve.

Under maternal inheritance, mitochondrial evolution is thought to be restricted to female lineages, leading to the accumulation of male-harming mutations when they have neutral or beneficial effects on female fitness (i.e. “mother’s curse”[26,28]). Initial evidence for the mother’s curse hypothesis (MCH) centered on evidence that sperm motility in humans and other taxa was linked to mitochondrial function[29,30,28] therefore sperm are more exposed to deleterious effects of mtDNA mutations than ova. The MCH has also been proposed to impact viability, and even to contribute to the taxonomically widespread pattern of shorter male lifespans[31]. Theoretical work, however, shows that conditions favoring the spread of mother’s curse mutations can be limited by even modest levels of inbreeding and kin selection, allowing for an adaptive response to mtDNA alleles impacting male viability and fertility[32–34]. Nevertheless, interest in MCH has increased steadily in recent years leading to formal tests of the hypothesis, mostly in *Drosophila* but also in plants (reviewed in [35]), utilizing carefully constructed mitochondrial-nuclear (mitonuclear) hybrids with divergent mtDNA haplotypes backcrossed into an isogenic nuclear background. The most direct test of MCH focuses on the effects of mutational variation in mtDNA, which are predicted to confer larger trait variance in males. A second major prediction focuses on the subset of alleles that are directly sexually antagonistic, which could lead to a negative intersexual genetic correlation across haplotypes. A third prediction is that experimental disruption of mitonuclear coadaptation will be more detrimental to males than females. Studies using the mitonuclear hybrid approach have shown both support[31,36–40], and rejection[41–44] of the mother’s curse hypothesis.

Understanding the pervasiveness of mother’s curse is crucial given its fundamental implications for the evolution of sexual dimorphism, as well as very practical consequences for conservation breeding programs and medicine (particularly mitochondrial replacement theory and male-biased mitochondrial disease). While past work has focused on *Drosophila*, the predominance of maternal inheritance of mitochondria in eukaryotes allows for a taxonomically diverse investigation of the generality of mother’s curse. Here we test predictions of the MCH using the splash-pool copepod *Tigriopus californicus*, an arthropod that shares a common ancestor with *Drosophila* around 409.3 – 536.6 million years ago[45]. We used a genetic panel of mitonuclear hybrids to assay both fertility and longevity. As fertility genes are likely sex-specific and viability genes are likely shared between the sexes (see review[46]), we predict that mitonuclear mismatch will lead to stronger sex bias for fertility than longevity. Additionally, we measured the impact of high environmental temperature on these traits, as mild mitonuclear mismatch can be expected to be unmasked under higher metabolic demand[47]. Populations of *T. californicus* show extreme genetic divergence over short geographical distances[48–50] while remaining somewhat tolerant of hybridization, producing viable and fertile offspring[51,52]. Nucleotide divergence in *T. californicus* extends across the mtDNA genome, involving coding and noncoding genes, and can exceed 20% between distant populations. *T. californicus* also differ from standard animal models in that they lack heteromorphic sex chromosomes - sex is determined by the inheritance of multiple, unlinked genes in combination with environmental effects (polygenic sex determination)[53–55]. While rarely utilized in evolutionary biology, polygenic sex determination provides an advantage for testing sex-specific mitochondrial effects as sex differences cannot be attributed to the effects of sex chromosomes. Importantly, despite the lack of heteromorphic sex chromosomes, there is marked sexual dimorphism in *T. californicus* and considerable evidence for sex-specific effects in multiple traits (thermal tolerance[56,57]; salinity tolerance[58,59]; resistance to toxins[59]; intrinsic hybridization stress[54]) in which males are generally the weaker sex.

## Methods

### Population sampling

Populations of *T. californicus* are distributed in supralittoral “splash pools” along the Pacific coast of North America. Copepods were originally collected from supralittoral pools at four locations (Figure 1A): Friday Harbor, WA (FHL: 48.55 N,123.01 W), La Jolla, CA (LJ: 32.83 N,117.27 W), Bird Rock, CA (BR: 32.82 N,117.27 W), and San Diego, CA (SD: 32.7460 N,117.2551 W). Isofemale lines were initiated from a single female fertilized by a single male, and maintained at 20°C on a 12h:12h light:dark cycle in standard culture media (three times filtered seawater from Wrigley Marine Science Center, Santa Catalina Island, CA; 0.1g/l of Spirulina and 0.1g/l ground Tetramin (Tetra®) Tropical fish food flakes). All lines were maintained in petri dishes for ≥ 5 years under these conditions and periodically fed additional Spirulina and ground Tetramin^®^ tropical fish food and rehydrated with DI water whenever necessary to maintain approximately 35 ppt salinity. Prior to experimentation, we sequenced samples from each isofemale line at the 12S ribosomal RNA locus (Forward primer: CCAAGATACTTTAGGGATAACAGC; Reverse primer: CTGTTCTATCAAAACAGCCCT; fragment size: 687 base pairs) to rule out cross-contamination.

**Figure 1.**
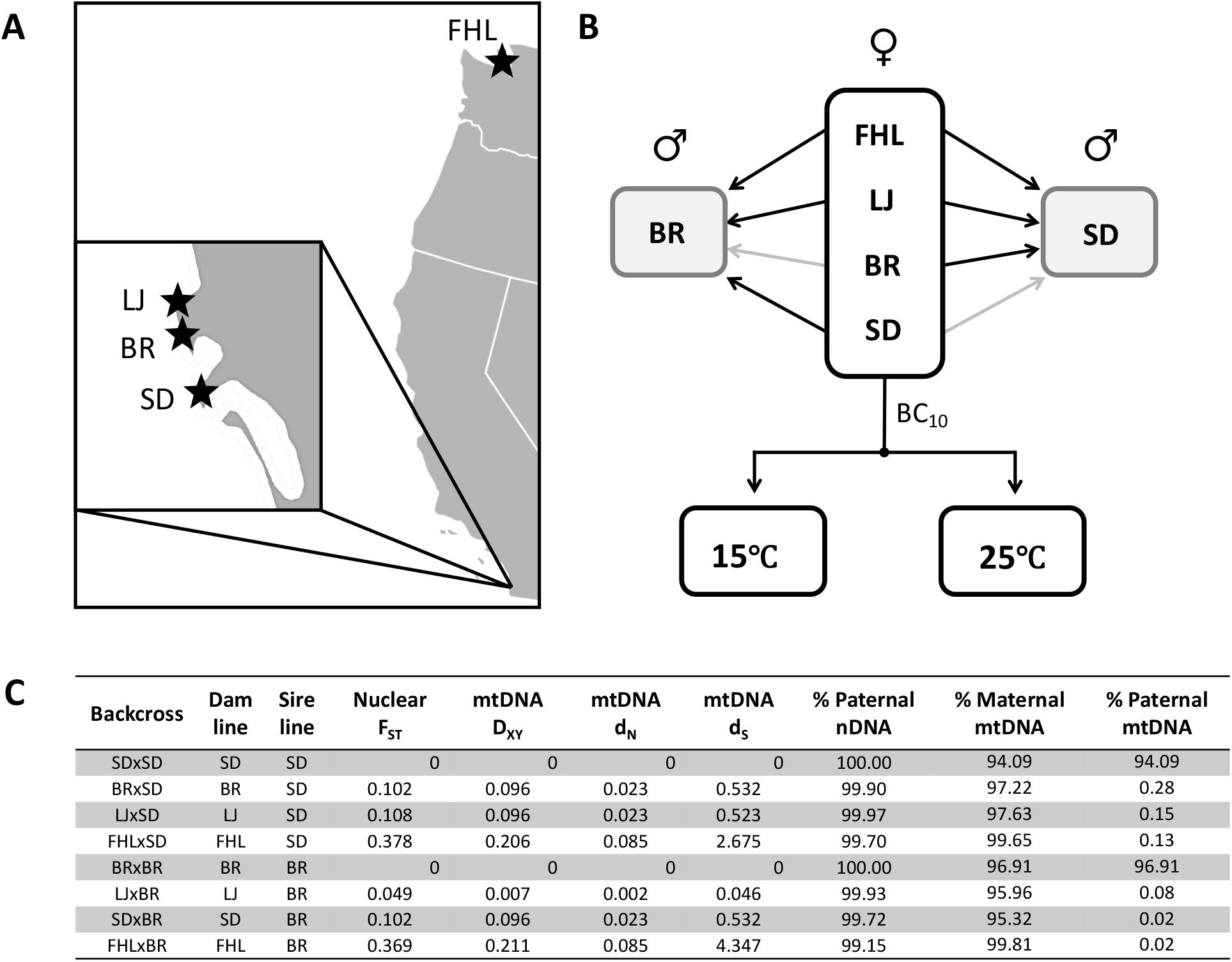
Construction of mitonuclear hybrid panel. (A) Geographic location of populations used in this study – SD (San Diego, CA), BR (Bird Rock, CA), LJ (La Jolla, CA), FHL (Friday Harbor, WA). (B) Experimental design. Dams from the central panel were backcrossed to sires from BR and SD for 10 generations. Grey arrows indicate experimental control lines. Individuals from each line were reared at two contrasting temperatures, 15°C and 25°C. (C) DNA sequencing results from pool-seq. Measures of divergence (F_ST_, D_XY_, d_N_, and d_S_) were calculated between dam and sire line. Percent paternal nDNA and percent maternal/paternal mtDNA indicate minimal contaminating nDNA remaining in backcrossed individuals and the absence of paternal leakage of mtDNA.

### Mitonuclear hybrid panel

In *Tigriopus*, meiosis is chiasmatic in males but achiasmatic in females, yielding recombinant sperm and non-recombinant ova. This allows us to backcross mitochondria from the population of the dam into that of the sire while preventing nuclear recombination between populations. Because sexes in *T. californicus* are not easily distinguished before maturity, we allowed males to clasp female siblings with their geniculate first antennae in order to ensure the sex of females. We moved clasped pairs onto filter paper and quickly separated them using entomological pins under a microscope. We backcrossed FHL, BR, LJ, and SD dams to SD and BR sires for 10 generations (Figure 1B), resulting in an average proportion of 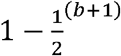 (where *b* is equal to the number of backcross generations) or 99.95% of the sire’s nuclear genome, assuming no selection. We chose the SD and BR lines for our isogenic nuclear backgrounds because SD was the source for the *<u>Tigriopus californicus</u>* genome assembly, BR is the genetically closest resequenced population to SD, and it would be experimentally intractable to perform a full factorial design. For each cross, we established 4 replicates, each maintained in 12-well plates containing standard culture media. All crosses were fed weekly with additional Spirulina and ground Tetramin^®^ fish food and rehydrated with DI water whenever necessary to maintain approximately 35 ppt salinity.

### Trait measurement and modeling

For each of our mitonuclear hybrid lines, we measured three fitness components (fertility, lifespan, and age-specific rate of survival) for males and females reared at two contrasting temperature regimes (15°C and 25°C). In order to achieve an adequate sample size for each mtDNA haplotype and sex, it was necessary to set up multiple families from each mitonuclear hybrid line. Each family was composed of up to four clutches of fertilized eggs derived from a single mitonuclear hybrid or control female (dam). Dams were maintained at standard conditions and checked daily for the presence of hatched eggsacs. For each dam, we alternated moving up to four hatched eggsacs to 15°C and 25°C (Figure 1B) so that clutch order was orthogonal to temperature in both fertility and longevity experiments. Individuals were maintained at the same temperature for the duration of the fertility or longevity experiment.

### Longevity

Individual animals were separated into wells in a 24-well plate once they developed to the juvenile copepodid stage and were checked three times a week. The first measurement we recorded was the sex-specific age of reproductive maturity. For males, this occurred with the development of geniculate first antennae (claspers) and for females with the development of eggs, which can be seen through the translucent body. We continued monitoring longevity experiments three times a week, recording the age of death of individuals until all had expired.

We performed linear mixed-effects modeling on family sex ratio using sex information from our longevity experiment, comparing mtDNA x nDNA interactions with the impact of mitonuclear status on family sex ratio. We also conducted a separate experiment, using different families, measuring mortality occurring before sexual maturity (Document S1, Figure S1). We were able to measure the naupliar, copepodid, and total development rate, and perform linear mixed-effects modeling investigating the influence of genetic (mtDNA, nDNA) and environmental (temperature) influence on these juvenile life-history parameters.

### Sex-specific fertility

For female fertility experiments, all individuals from hatched eggsacs were separated into individual wells in a 24-well plate once they developed to the juvenile copepodid stage. To each well, we added a single wild-caught adult male from the appropriate locality (SD or BR), matching the paternal line of the individual. Males were added to every well because sexes are difficult to distinguish morphologically in immature *T. californicus* and we wanted to ensure that each female was paired with an adult male before sexual maturity. Mate clasping typically occurs between adult males and juvenile females, and rarely occurs with mature females once they begin extruding non-fertilized eggsacs. We also ran experiments utilizing sib-pairs in parallel to compare the impact of mating with mitonuclear hybrid males versus wild-type males on female fertility (Document S1). Plates were maintained at the appropriate temperature (15°C or 25°C) and checked three times a week until females produced an eggsac. As we utilized several hundred plates in our experiment, it became necessary to maximize incubator space. Therefore, we placed gravid females from the same family and clutch together in petri dishes until they reached experimental age. Because individuals develop twice as fast at 25°C as at 15°C, we used temperature-specific ages for our experiment in order to minimize age-related variation. Once individuals reared at 25□ reached 35 days of age, and individuals reared at 15□ reached 70 days of age, we moved them into separate dishes and maintained them at the correct temperature, checking them three times a week until eggsacs hatched. Upon hatching, each individual larval nauplius was counted under a dissecting microscope to measure female hatching number.

For male fertility experiments, we removed males from each of the four clutches in a family and allowed them to age for the appropriate temperature (35 days old for 25°C and 70 days old for 15°C). Once at the appropriate age, each male was pipetted into a petri dish with six wild-caught juvenile females from the appropriate locality (BR or SD) to match the paternal nuclear line. To ensure that wild-caught juveniles were female, we selected clasped pairs from culture within 0-2 weeks of collection and separated pairs under a dissecting microscope. Care was taken to provide males with females of consistent and reasonably large size, to reduce variation across experiments. Each experimental male and six wild-caught females were allowed 14 days to mate in a petri dish at the appropriate temperature. After 14 days, females were isolated into a 6-well plate and scored for the presence or absence of any offspring indicating a successful mating. We recorded male remating success as the ratio of successful matings to the number of surviving females.

### Parametric and nonparametric trait modeling

For our trait modeling, we performed three types of linear models: sex-effect modeling, coevolutionary status modeling, and best-fit models. Sex-effect models measure sex effects impacted by mtDNA and nDNA variation and their interactions. If mother’s curse mutations are present, they may exist in additive (Sex x mtDNA) or epistatic (Sex x mtDNA x nDNA) form. Sex differences may also be encoded by nuclear genes, described by the (Sex x nDNA) term. In order to compare fertility experiments across the two sexes, individual measurements of fertility were normalized to the mean of the appropriate matched control line reared at the same temperature (e.x. male *i* from FHLxSD at 15□ / mean of all males from SDxSD at 15□). We performed model testing on the four sex-effect models using the *model.sel* function in the MuMIn[60] R package, which ranks models based on their AIC_C_ values. Model contrasts for sex differences were performed using the *pairs* function from the emmeans[61] R package. Coevolutionary status models measure the impact of disrupting coevolved mitonuclear combinations on sex-specific data grouped by mtDNA haplotype. These models included temperature and coevolutionary status as fixed-effect terms, compared mitonuclear hybrids of both nDNA backgrounds with both matched controls and included nDNA as a random term. Mitonuclear hybrids with the SD or BR mtDNA were only compared to one matched control. Modeling for best-fit utilized an “all-subset” approach, where all possible combinations of predictors were compared using the *MuMIn* function *dredge*. Models with the highest weight and lowest AICC were designated as best-fit. Both fertility and longevity models included temperature, nDNA, mtDNA and all possible interactions as fixed-effects terms in initial models and were performed separately for each sex. For fertility, we used raw sex-specific fertility (males: remating success, females: hatching number). All linear mixed-effects modeling was performed using the *lmer* function from the lme4[62] R package and included clutch nested within family as a random-effect term. We used type III ANOVA to estimate significance for model main effects and interaction terms using the *Anova* function from the car[63] R package.

We performed parametric regression of survivorship for each nuclear, temperature, and sex combination against the Gompertz distribution [64], estimating the shape (α) and rate (β) parameters using the R package flexsurv[65]. Parameters were estimated by maximum likelihood, with those defined to be positive estimated on a log scale. Confidence intervals were estimated from the Hessian at maximum, and then transformed back to the original scale (days). The shape of the resulting distributions, ln(α), describe the initial mortality rate or frailty of the samples, while the rate (β) parameter describes the age-independent rate of mortality acceleration or rate of senescence. We employed Kruskal-Wallis one-way analysis of variance to estimate the significance of parameter estimates produced by parametric survival models using the *kruskal.test* function from the stats[66] R package.

### Mitochondrial coefficient of variation

One prediction of the mother’s curse hypothesis is that the accumulation of male-affecting variation should lead to a higher mitochondrial coefficient of variation (*CV_mt_*) in males than in females. Using the *boot*[67,68] package in R, bootstrapped *CV_mt_* was computed by resampling raw trait means for each mitochondrial line using 1000 replicates with replacement and calculating the coefficient of variation among mitochondrial lines (*CV = σ/μ*) for each nuclear, sex, and temperature combination. Hypothesis testing was performed to compare male and female *CV_mt_* using a sign test as implemented in the *SIGN.test* function in the BSDA[69] R package.

### Genetic correlation among haplotypes

Mother’s curse mutations are likely to reach fixation fastest when they confer a fitness advantage to females, resulting in sexual conflict when they simultaneously result in low fitness in males. When enough mtDNA variation is involved in sexual conflict, a negative intersexual correlation may be observed across haplotypes for a given trait. We measured the correlation between haplotype means for fertility and median longevity, comparing traits within sexes (intrasexual correlation) and also comparing the sexes for a given trait (intersexual correlation). We used bootstrapped estimates of Pearson’s coefficient using the boot R package along with the *cor* function from the stats R package, recording 1000 samples for each comparison. Equi-tailed, two-sided nonparametric confidence intervals were generated from the first order normal approximation with a confidence level of 0.95 using the *boot.ci* function from the boot package in R.

### Whole genome sequencing

For each isofemale and mitonuclear hybrid line, we extracted DNA from 10 males and 10 females using Zymo quick-DNA preps and quantified DNA yields using Qubit HS dsDNA quantification kits. Because female *T. californicus* yield approximately 2.12x more DNA than males, we pooled 2 ng of DNA from each of the 20 individuals per line for library preparation. DNA pools were sheared using a Covaris S2 sonicator with microTUBE-50 AFA tubes, and sequencing libraries were prepared using Qiagen QIAseq Ultralow Input kits following instructions, using 8 PCR cycles during library amplification. Mitonuclear hybrid libraries were pooled together in equimolar amounts and sequenced together on a single lane of Illumina HiSeq4000 (PE-150), and the parental isofemale libraries were pooled and sequenced on a separate lane. Sequence reads were quality trimmed using trimmomatic[70] v0.36, and paired reads were filtered for mitochondrial reads using the filter_reads.pl perl script included with the NOVOPlasty[71] software. We assembled consensus mtDNA genomes from each sequenced isofemale line, by utilizing the appropriate mtDNA reference from a previous publication which included the SD, BR, and FHL populations[23]. A reference mtDNA genome did not exist for the LJ population, so those reads were mapped to the BR mtDNA reference. We annotated our *de novo* assembled genomes using the Mitos[72] 2.0 software, each containing all 13 protein coding genes, and the majority of non-coding RNAs. Following this, we mapped all trimmed and paired reads to the *T. californicus* reference genome assembly (NCBI accession: GCA_007210705.1) along with the appropriate mitochondrial genome using stampy[73] v1.0.32 (substitutionrate=0.05). Following the removal of PCR and optical duplicates using picard[74] MarkDuplicates, we called variants for each sequenced pool using samtools[75] mpileup.

We estimated pairwise nuclear divergence for isofemale lines in a sliding window approach. For each comparison, we calculated F_ST_ using a sliding window of 10,000 bp, requiring a depth of 40x coverage, and minimum PHRED base quality score of 30 using the fst-sliding.pl perl script included with popoolation v1.2.2. For parental mitochondrial divergence, we aligned whole mtDNA genomes using progressiveMauve[76] along with mtDNA genome sequences from an outgroup population from San Roque, Baja California Sur, Mexico to produce rooted trees, and measured divergence (*d_XY_*) using the calculate-dxy.pl perl script included with popoolation v1.2.2. We also measured divergence at synonymous and nonsynonymous sites by aligning protein-coding sequences with MACSE [77] v2, constructing neighbor joining trees with ClustalW [78] and analyzing these data with codeml from the PAML suite [79] using the null model (M0), assuming a uniform selective pressure among sites.

### Confirming mitonuclear genetic status of hybrids

To confirm the isogenic nuclear status of our lines, we measured allele frequencies at a set of divergent sites for each paternal line’s nuclear genome. A Fisher’s exact test for significant allele frequency differences for nDNA from the maternal and paternal lines was conducted using popoolation[80] v1.201, requiring a minimum depth of 20x coverage. In mitonuclear hybrids, sites with no significant differences (alpha > 0.05) to the paternal line were considered “paternally concordant” and significantly different sites (alpha <= 0.05) were considered “paternally discordant”. Because significance may be determined by variations in the same alleles between lines, we recorded paternally discordant sites with alleles matching maternal lines as “maternally concordant” when they were absent in the paternal nDNA. Paternal concordance rates were recorded by measuring the proportion of sites that were not maternally concordant to the total number of sites. Lee and Willett[81] recently reported the presence of paternally inherited mtDNA, likely due to paternal leakage, in F1 hybrid *T. californicus*. This paternal mtDNA was found in a high frequency of individuals (up to 59%) but a very low frequency of reads (<0.01% in pooled samples). We therefore tested for paternally inherited mtDNA in our mitonuclear hybrid lines. This was accomplished by mapping trimmed and paired sequence reads to the reference genome as well as all mtDNA haplotypes simultaneously. We measured depth of coverage for each mtDNA haplotype using samtools depth command to identify any potential sources of contamination.

## Results

### DNA sequencing

Illumina HiSeq 4000 sequencing yielded an average of 34.7 million reads for mitonuclear hybrid line pools, and 76.1 million reads for inbred line pools. The read sequences were deposited at the National Center for Biotechnology Information (NCBI) Sequence Read Archive (SRA) under the project accession number PRJNA840866. Following quality trimming and mapping of paired reads to the appropriate references, an average of 29 million reads mapped for cybrid pools, and 61 million reads mapped for maternal and paternal line pools. Average nuclear genome coverage across inbred lines was 91.88% at a mean depth of coverage of 26x. Across the mitochondrial genome, the mean depth of coverage was 2670X, and genome coverage was 100%. Nuclear divergence (F_ST_) between paternal lines ranged from 0.049 to 0.369 in lines with BR nDNA, and from 0.102 to 0.378 in lines with SD nDNA (Figure 1C). Mitochondrial divergence (d_XY_) between parental lines ranged from 0.7% to 21.1% in BR nDNA lines, and 9.6% to 20.6% in SD nDNA lines. The mitonuclear hybrid panel represented a wide range of mitochondrial variation, with as few as 13 amino acid substitutions between the BR and LJ mitotypes, 170 aa substitutions between the BR and SD nuclear line mitotypes, and 555 aa substitutions between SD and FHL haplotypes. The highly divergent FHL haplotype also differed from the BR & LJ haplotypes by 546 aa substitutions.

For each mitonuclear hybrid line, we measured allele frequencies at an average of 580k divergent nDNA SNPs between parental lines. We found that alleles were mostly concordant with paternal nDNA, with limited evidence of introgression from maternal nDNA, with paternal concordance rates ranging from 99.15% - 100% (Figure 1C). To test for potential mtDNA contamination via paternal leakage or experimental error, we compared depth of coverage across all mtDNA haplotypes, since mtDNA positions and alleles are not comparable across haplotypes. We found that the percent maternally matching depth ranged from 94.09% to 99.81%, while the percent of paternally matching depth ranged from 0.02% to 0.28% providing limited evidence for contaminating mtDNA (Figure 1C). While we cannot determine with certainty whether this result confirms paternal leakage, it is important to note that paternally matching depth measured in our study may also be confounded with nuclear-encoded mitochondrial DNA fragments (numts) which also match the paternal mtDNA haplotype. Further, because pooled-DNA sequencing lacks haplotype information, we are also unable to differentiate such low-frequency variants from sequencing error (0.1-1%).

### Trait measurements and modeling

Despite initially setting up four replicates for each mitonuclear hybrid line, low productivity led to extinction of several replicates, forcing us to proceed with one replicate per line to keep our experiment balanced. We measured the hatching number for 182 females coming from 41 families reared at 15°C and 216 females from 40 families reared at 25°C. Male remating success was measured for 491 males, using a total of 2,458 wild-caught females. Since 378 females died over the course of the experiment, we discarded 31 males that had fewer than four out of six females remaining in order to reduce variation associated with differential access to females. Therefore, our male remating experiment is based on 235 males from 68 families reared at 15°C and 225 males from 76 families reared at 25°C. The average male successfully copulated with 2.45 females in the span of 14 days when reared at 15□, declining to 2.04 females at 25□ representing a 17% reduction. In the average female clutch, 30.05 eggs hatched at 15□, declining to 21.50 eggs hatching at 25□ representing a 28% reduction. Female hatching number declined an average of 31.16% in mitonuclear hybrid lines when females were bred with siblings as opposed to wild-caught males (Document S1, Figure S1). Lifespan, frailty, and rate of senescence was measured for 3,072 individuals, with 1,531 individuals from 89 families reared at 15°C, and 1,541 individuals from 77 families reared at 25°C.

Experiments on early developmental stages indicate that mortality rate occurring before sexual maturity was very high, ranging from 14.6% to 90.6% mortality depending on cross and temperature (Document S1, Figure S2). Naupliar stages had the highest mortality, ranging from 4.2% to 84.4%, while copepodid stages ranged from 1.5% to 27.9% mortality. Linear mixed-effects modeling showed that measurements describing the development rate varied significantly with mtDNA x nDNA interactions and rearing temperature, with rates increasing at higher temperature.

### Sex-effect modeling

Model contrasts revealed mostly male-biased values for longevity (longer lifespan), and mixed bias for fertility with three out of six significant contrasts being female biased (Figure 2A). Model selection comparing the four sex-effects models (Table S1) revealed that models including three-way sex x nDNA x mtDNA interaction term fit the trait data best in all temperature and trait combinations except for relative fertility at 15□, which was better fit with sex x nDNA interactions (ΔAIC = 17.35). None of the sex-effects models uncovered significant sex x mtDNA interactions (Table S2). However, we do see that sex x nDNA and sex x mtDNA x nDNA interactions are a significant influence on fertility variation (Figure 2B).

**Figure 2.**
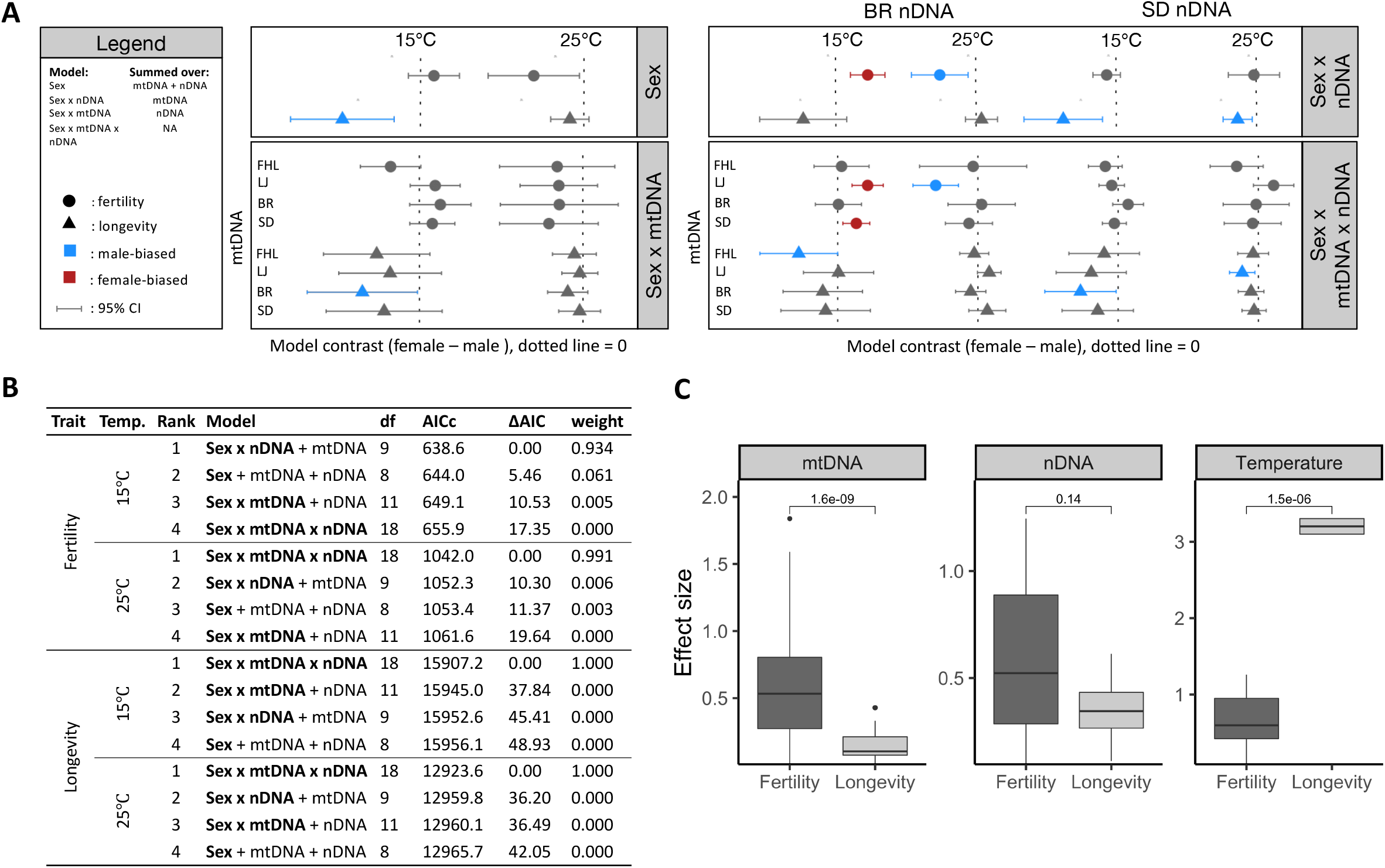
Trait modeling results. (**A**) Model contrasts between the sexes for the four sex-effect models. Models were performed separately for each temperature. Model contrasts for the “Sex” model were summed over all mtDNA and nDNA sources, “Sex x nDNA” contrasts were summed over mtDNA, and “Sex x mtDNA” contrasts were summed over nDNA. Circles represent sex contrasts for fertility and triangles represent longevity. The vertical dotted line represents contrasts (female – male) = 0.0, indicating identical estimated marginal means. Contrasts with 95% confidence intervals that fall on the left side are negative and thus significantly male-biased and coloured blue, while those falling on the right side are positive, female-biased, and coloured red. (**B**) Sex-effect model type III ANOVA table. Each row represents a separate model, with columns representing modeled fixed effects. Cells are marked with a dash when a variable is not included in the model. Abbreviations: NS – non significant (p <0.05), * – p ≤ 0.05, ** – p ≤ 0.01, ***– p ≤ 0.001. (**C**)Genetic and temperature effect size comparisons between fitness traits. We estimated Cohen’s *d* for all pairwise contrasts among temperature, mtDNA, and nDNA groupings. Contrasts were made on best fit linear mixed effects models for each sex and trait grouping (sex-specific fertility, longevity). Significance was determined using two-sample unpaired Wilcoxon rank sum tests with a two-sided alternate hypothesis and 95% confidence intervals. Sample sizes for contrasts were mtDNA, n = 96; nDNA, n = 32; and Temperature, n = 32.

### Mitonuclear coevolutionary status modeling

Fertility was negatively impacted by mitonuclear status (matched vs. mismatched), but only in females possessing the FHL mtDNA haplotype (females: *χ*^2^ = 17.90, p_Bonf._ = 7.2e-05; males: *χ*^2^ = 0.13, p_Bonf._ = 1.0), while no other trait was significant using the Bonferroni-corrected cutoff (Table S3). The estimated marginal mean for matched female hatching number was 26.9 offspring, while for mismatched FHL females it was 16.5 offspring. Females with FHL mtDNA also showed a trend of delayed development at the naupliar (females: *χ*^2^ =5.20, p_Bonf._ = 0.0675; males: *χ*^2^ = 1.16, p_Bonf_ = 1.0) and copepodid (females: *χ*^2^ =4.39, p_Bonf_ = 0.1086; males: *χ*^2^ =1.52, p_Bonf_ = 1.0) stages, although these values were less than alpha = 0.05 and not the Bonferroni-corrected alpha. The estimated marginal means for naupliar and copepodid development rate in matched females were 8.68 and 12.1 days respectively, while the means for mismatched FHL females were 12.08 and 14.9 days. Family sex ratio was significantly influenced by mitonuclear status *χ*^2^= 18.51, *p* = 1.7e-5), as indicated in linear mixed-effects modeling (Document S1, Figure S3), as well as nDNA x mitonuclear status interactions *χ*^2^= 28.77, *p* = 8.1e-8). Results using mortality-corrected family sex ratio data were also consistent with uncorrected family sex ratio. (Table S4).

### Best-fit models

Temperature significantly impacted all traits by reducing their average values (Table S5). Notably, mean longevity was reduced by 102 days in 25°C, an almost 10 day decrease per degree reduction. Significant mtDNA effects were only seen in females for both hatching number (*χ*^2^= 20.36, *p* = 1.43e-4) and longevity *χ*^2^= 14.66, *p* = 0.0021). Variation in nDNA impacted longevity in both sexes (males: *χ*^2^= 20.36, p = 6.42e-6; females: *χ*^2^= 8.31, p = 0.0040) but neither measure of fertility (female hatching number: *χ* = 1.37, *p* = 0.2416; male remating success: *χ*^2^= 0.15, *p* = 0.6991). Additionally, effects of nDNA were contingent upon rearing temperature, with significant interactions between the two terms for male remating success (*χ*^2^=7.41, *p*= 0.0065), male longevity (*χ*^2^= 4.19, *p* = 0.0407) and female longevity *χ*^2^= 4.05, *p* = 0.0442). Interactions between mtDNA and nDNA were also significant for longevity in females *χ*^2^= 12.03, *p* = 0.0073) and in a three-way interaction with temperature for male remating success *χ*^2^= 16.95, *p* = 7.22e-4). Mitochondrial effects impacted fertility more than longevity (Figure 2C, *p* = 3.2e-4), while nuclear effects impacted both traits equally (*p* = 0.95) and temperature affected longevity more than fertility (*p* = 1.5e-6). Linear mixed-effects modeling of family sex ratio indicated significant influence of temperature (*χ^2^* = 10.02, *p* = 0.0015), mtDNA haplotype (*χ^2^* = 28.35, *p* = 3.1e-6), as well as interactions between mtDNA and nDNA (*χ*^2^= 31.98, *p* =1.6e-8) and temperature and nDNA *χ*^2^= 29.94, *p* = 1.4e-6; Document S1, Figure S3).

### Parametric survivorship modeling

Survivorship curves indicate a drastic reduction in longevity shifting from 15□ to 25□, with SD nDNA individuals having higher survivorship than BR nDNA individuals at 25□ (Figure 3A). Additionally, we see that median longevity is sex-biased in SD nDNA individuals reared at 15□, with higher male longevity (Figure 3A insert, Wilcoxon rank sum test, p = 0.004). Rearing temperature affected both frailty *χ*^2^= 22.55, *p* = 2.05e-6) and rate of senescence *χ*^2^= 23.27, *p* = 1.41e-6) in a similar manner, with the higher temperature leading to increased frailty and increased rate of senescence (Table S6). Mitochondrial haplotype showed no significant contribution to Gompertz parameters when reared at either 15°C or 25°C (Table S6). We found significant sexual dimorphism in frailty and senescence in individuals reared at 25°C, but not those reared at 15°C (Figure 3B). When reared at 25°C, females had lower frailty than males, but a higher rate of senescence and while this pattern held in both nDNA backgrounds, it was nonsignificant in BR nDNA lines (Table S6).

**Figure 3.**
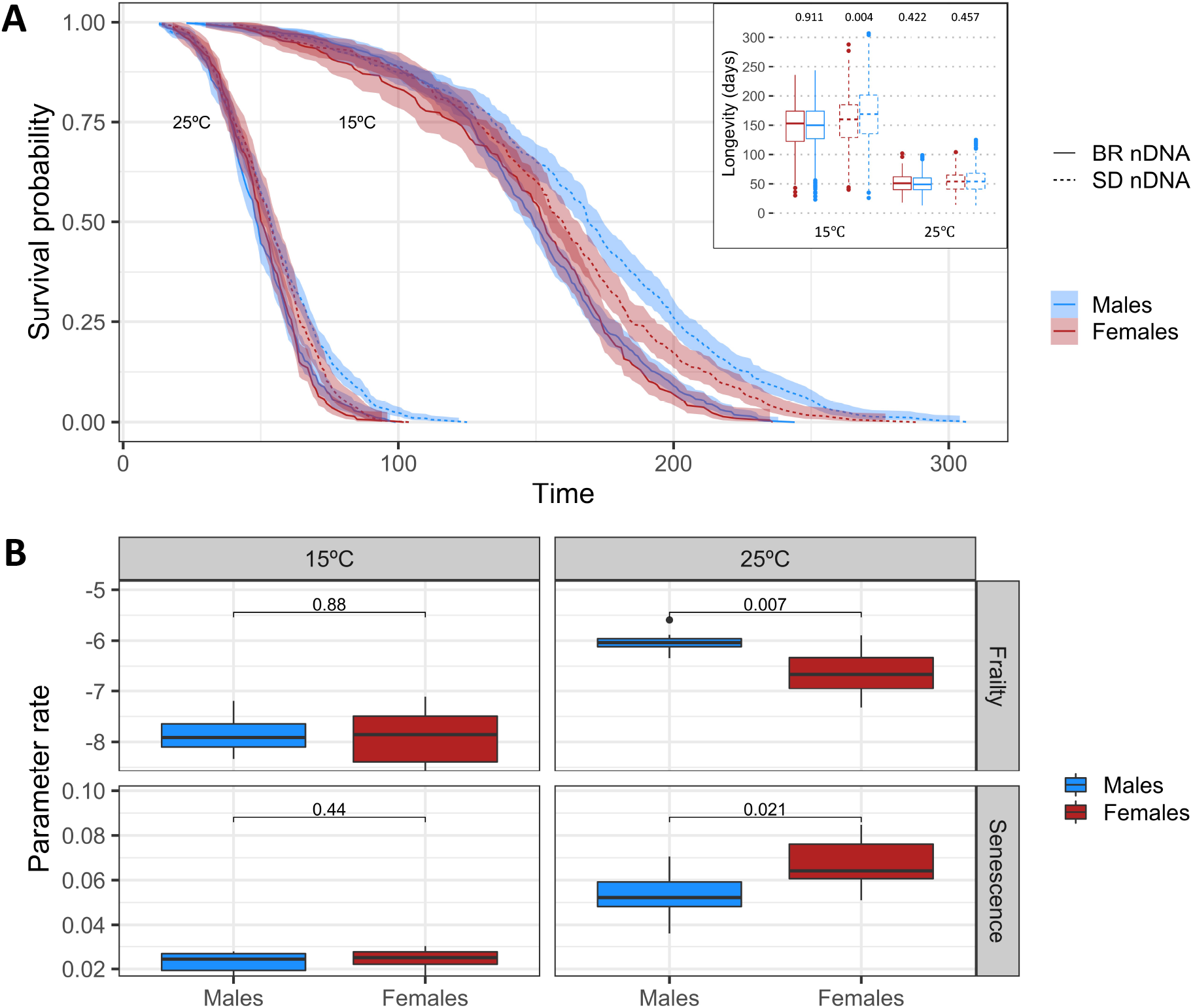
Males outlive females and have lower rates of senescence and higher frailty. (A) Sex-specific survivorship curves based on nDNA and temperature groupings. Median longevity shown in insert with p-values indicating results of two-sample unpaired Wilcoxon rank sum tests with a two-sided alternate hypothesis and 95% confidence intervals. (B) Sex contrasts for Gompertz parameters. We performed Wilcoxon tests as above to identify parametric differences in the rate of aging between the sexes.

### Mitochondrial coefficient of variation

Females had significantly higher *CV_mt_* than males in seven out of eight experimental groupings (Figure 4A, Table S7). The median difference between male and female coefficients in these groupings ranged from 0.0039 to 0.1061. In contrast, fertility in BR nDNA individuals at 25°C showed higher male *CV_mt_* than females (median difference = −0.0993, CI = (−0.11, −0.09), *p* < 2.2e-16).

**Figure 4.**
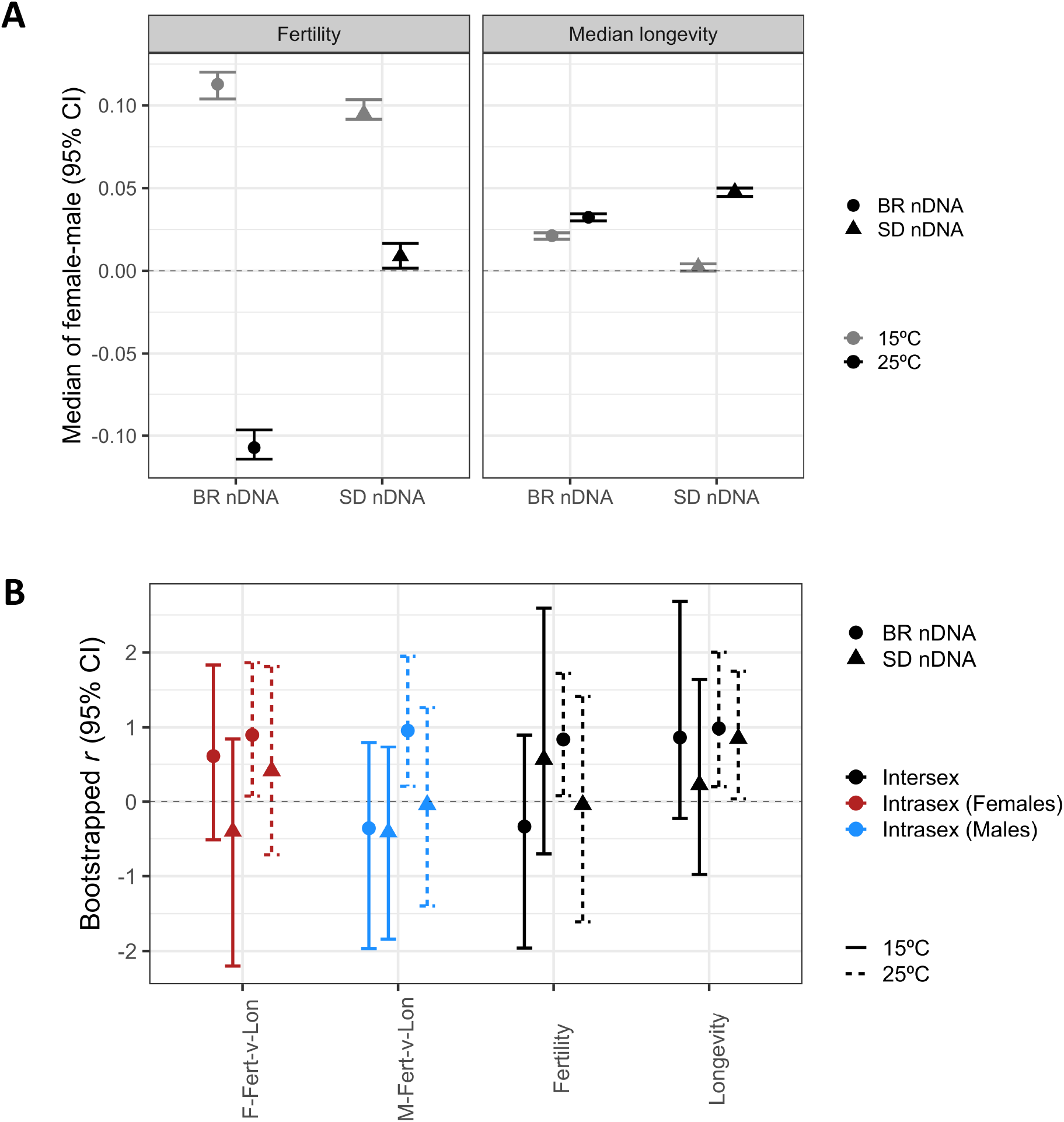
Testing the mother’s curse hypothesis with mitonuclear hybrid *T. californicus*. (A) Sex differences in mitochondrial coefficient of variation (CV_mt_) for fertility and longevity. We estimated CV_mt_ for both fertility and median longevity by temperature and nuclear line and performed a two-sided sign test on bootstrapped estimates (1000 samples). The sign test compares the difference between female and male median values with negative values indicating higher male CV_mt_, while positive values indicate that females have higher CV_mt_.. The majority of groupings indicate higher mitochondrial genetic variation in females in contradiction to the mother’s curse hypothesis. Only fertility in BR nDNA individuals reared at 15°C had significantly higher mitochondrial genetic variation in males. (B) Intra- and intersexual genetic correlation for female and male fertility and longevity across mitochondrial haplotypes. Bootstrapped Pearson’s correlation coefficients (95% confidence intervals) were plotted for both rearing temperatures (15°C: solid lines, 25°C: dotted lines). Abbreviations: F (females), M (males), fertility (females: hatching number, males: remating success), longevity (mean longevity). All significant intersexual correlations indicate positive values, suggesting the absence of sexually antagonistic pleiotropy predicted by the mother’s curse hypothesis.

### Genetic correlation among haplotypes

Intersexual correlations for median longevity (*r* = 0.99; CI_95_ = 0.20, 2.00) and fertility (*r* = 0.84; CI_95_ = 0.08, 1.72) were significantly positive in BR nDNA individuals reared at 25°C, while correlations for median longevity in SD nDNA individuals were also significantly positive (*r* = 0.85; CI_95_ = 0.04, 1.75) in individuals reared at 25°C (Figure 4B). Intrasexual correlations between fertility and longevity were significantly positive in BR nDNA individuals of both sexes reared at 25°C (males: *r* = 0.96; CI_95_ = 0.21, 1.95; females: *r* = 0.90; CI_95_ = 0.08, 1.86). No measures of intersexual or intrasexual correlation were significantly negative.

## Discussion

Our study sought to explicitly test three predictions arising from the mother’s curse hypothesis: higher mitochondrial genetic variance in males for fitness traits, negative intersexual correlation among mitochondrial haplotypes for fitness traits, and higher impact of experimental disruption of putatively coevolved mitonuclear genotypes on males. Our results did not support these predictions. Females had a higher level of mitochondrial genetic variance in seven experimental groupings and lower variance in one grouping (fertility in individuals with BR nDNA reared at 25°C). We also found no evidence for sexual antagonism for fertility or longevity, with negative intersexual correlations found in two of eight experimental groupings (fertility in BR nDNA at 15□ and SD nDNA at 25□) having upper confidence intervals that extended well into positive ranges (0.90 and 1.41, respectively). Overall, the intersexual comparisons mimicked the intrasexual comparisons. Lastly, we found a significant negative influence of coevolutionary status on female fertility in hybrids with the FHL mtDNA, but not for male fertility contrary to the predictions of the mother’s curse hypothesis. Female hybrids with FHL mtDNA also show a trend of delayed larval development, while males do not. The FHL mtDNA is the most divergent haplotype (≥ 20.6% nucleotide divergence) while the remaining haplotypes are 0.7 - 9.6% divergence).

Detecting sex-bias in mitochondrial coefficient of variation for fitness traits is arguably the most direct test of the hypothesis since it emphasizes the differential accumulation of mutational variation between the sexes[82]. According to the mother’s curse hypothesis, mitochondrial mutations causing male infertility that are neutral in females are more likely to spread in a population through random drift since males are a dead-end for natural selection on mtDNA. However, this argument is best applied to randomly mating populations with no inbreeding. In populations with even modest levels of inbreeding, such as those with geographical population structure, females will also suffer fitness insults from mother’s curse mutations to a certain degree since they are more likely to mate with sub-fertile siblings[32,33,83]. The degree to which male subfertility influences female fertility fitness can also influence these dynamics. In monandrous species like *Tigriopus*, females mate only once[84,85], resulting in a direct relationship between male fertility and female offspring production. In this study, we showed that female hatching number dropped substantially when mated with siblings compared to wild-caught males, indicating the influence of male subfertility on female fertility. However, in mating systems where females mate with several males (e.g. *Drosophila*), females can be safeguarded against male infertility or subfertility[86]. As a result, selection to purge male-harming mtDNA mutations is expected to be strongest in species with monandry and weakest with polyandry whenever inbreeding is present[83]. Studies show that *T. californicus* avoid close inbreeding[87], but live in highly isolated, ephemeral populations making inbreeding all but inevitable[88,89]. Additionally, while inbreeding can limit mother’s curse by decreasing the conditions under which male-harming alleles can spread, it can also result in the spread of female-harming alleles that benefit males and lead to female-biased fitness variance [27,32]. The widespread female-biased fitness variance observed in our fertility and longevity experiments may therefore indicate the impact of historical inbreeding on the fixation of mitochondrial divergence sampled in our study.

Another factor that has been proposed to limit the spread of male-harming mtDNA mutations is paternal leakage of mtDNA, although we do not believe it played a major role in our results. We found a very low level of mtDNA reads of potential paternal origin (0.02% to 0.28%), and these could not be cleanly distinguished from numts or sequencing error. Haploid models of cytoplasmic inheritance suggest that small amounts of paternal leakage are not sufficient to rescue mother’s curse[90], with indirect selection on human males via inbreeding estimated to be up to 370 times stronger than direct selection on sperm performance due to paternal leakage [32]. Indeed, low levels of paternal mtDNA leakage do not appear to reliably cure mother’s curse, since levels of leakage higher than those reported in *Tigriopus* [80] are common in *Drosophila* crosses[91,92], where the strongest (albeit inconsistent) support for mother’s curse has been found.

We predicted that sex differences due to mitochondrial haplotype effects are greater for fertility than longevity, based on the expectation that fertility genes are more sex-specific than viability genes[93,94]. We also predicted that the consequences of mitonuclear mismatch should increase with metabolic rate associated with increased temperature. Efficient energy production depends on mitochondrial function, including mitonuclear coordination. Mild mitonuclear mismatches may be tolerated under low aerobic demand and yet become lethal when energetic demand become higher ^40,but see 82^. Increased temperature reduced measures of both fertility and longevity but was more dramatic with the latter. We uncovered significant mitonuclear effects impacting longevity at 25□ but not 15□. Additionally, we found that mitonuclear x sex effects were significant for fertility at 25□, but not 15□. Mitochondrial haplotype effects in our experiment impacted fertility more than longevity, however experiments on juvenile stages indicate a substantial amount of mortality, influenced by temperature and mitonuclear interactions, occurring before sexual maturity. Because *T. californicus* sperm are amotile, declines in male fertility are not attributed to loss of sperm motility – a key marker of male sterility in organisms with flagellated, motile sperm. Instead, declines in male fertility in our study are likely due to the number of mature spermatids in a spermatophore, the ability of sperm to maintain homeostasis while residing in the female reproductive tract, or behavior. High juvenile mortality may also impact our study as it could produce indirect selection on the adult traits under study, although any juvenile mortality precludes analysis of sex differences. Mortality occurring during the juvenile stages could not be assigned to either sex, as it is difficult to distinguish the sex of *Tigriopus californicus* juveniles before sexual maturity. However, while family sex ratio differed among crosses and by temperature, most crosses indicate male-biased sex ratios, which may indicate either increased mortality of females during the larval stages or the misexpression of sex determining loci in mitonuclear hybrids.

Survivorship analysis indicates that mtDNA variation had an inverse impact on frailty and senescence between the sexes, similar to the ‘Strehler-Mildvan’ (S-M) relationship[96]. When reared at 25°C, females had lower frailty than males while also having a higher rate of senescence, despite having similar median longevity. One explanation for this relationship is that the preferential survival of the most-frail female *T. californicus* may lead to an increase in the rate of senescence for females, since frail individuals are more susceptible to aging. Male *T. californicus* typically have lower tolerance than females to a variety of stressors, including temperature, and a greater transcriptomic response to oxidative stress[56,57,59,97,98].

## Conclusion

Like previous work[99,100], we found that the functional mtDNA effects were highly context-dependent, being influenced by the nuclear-background in which they were expressed, the temperature in which individuals developed, and/or the sex of the individual. We explicitly sought to test the main predictions of the mother’s curse hypothesis, finding patterns opposite to those predictions. Support for the mother’s curse hypothesis has been highly mixed to date, both theoretically and empirically, with many studies rejecting these predictions. While many studies directly testing predictions of the mother’s curse hypothesis have used *Drosophila*, the predominance of maternal inheritance of mitochondria in eukaryotes allows for a taxonomically diverse investigation of the generality of mother’s curse. Particularly, diversity in mating system should be considered for future studies, as the fitness of females that mate with only one male will be strongly determined by the fitness of her mate when there is inbreeding. Furthermore, sperm diversity (motile/amotile, storage/remating) should be highlighted in order to address the relevance of various mutational insults to male fitness potentially harbored by mitochondrial DNA. In light of the evidence presented here and elsewhere, we suggest that accumulation of male-harming mitochondrial mutations may not be a generalizable phenomenon, despite the prevalence of maternal mitochondrial transmission across eukaryotes.

## Acknowledgments

We are thankful to MacKenzie Partridge, Manuel Morales and Mi Rae Park for their help during the experiments and population maintenance. We also thank Ning Li and Scott Applebaum for their constructive criticisms and suggestions, which greatly improved the quality of the manuscript. Comments provided by anonymous reviewers also aided in improving the manuscript. The experiments reported here were supported by a grant (#1656048) from the National Science Foundation, Division of Environmental Biology, awarded to SE.

## SUPPLEMENTAL INFORMATION

Supplemental Information includes one document, three figures, and seven tables and can be found online.

