## Supplementary material for "Mitochondrial effects on fertility and longevity in *Tigriopus californicus* contradict predictions of the mother’s curse hypothesis": Figure S1

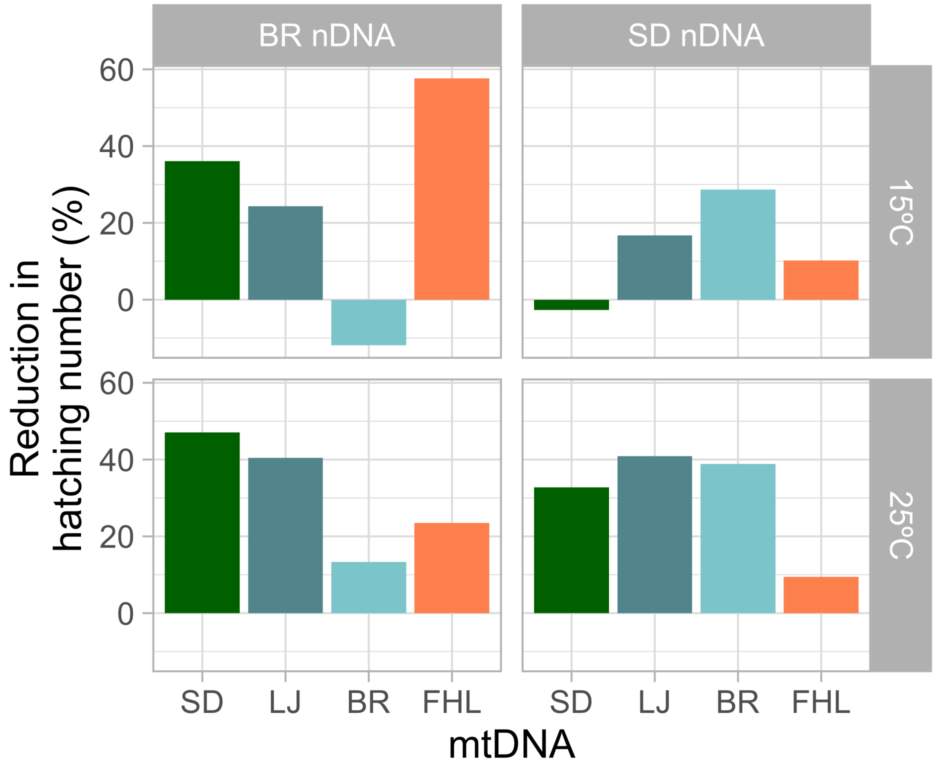


**Figure S1. Effects of inbreeding on female hatching number.** We conducted a separate female fertility experiment utilizing sibling crosses and compared the average hatching number for each experimental grouping with our outcrossed fertility experiment described in the main text. Inbreeding reduced hatching number in all crosses except for the matched control lines reared at 15℃, which increased. For the mitonuclear hybrid lines, inbred crosses experienced a 12.5 – 57.65% reduction in hatching number compared to outbred crosses suggesting the impact of male quality on female fitness.
