## Supplementary material for "Mitochondrial effects on fertility and longevity in *Tigriopus californicus* contradict predictions of the mother’s curse hypothesis": Figure S2

**A**

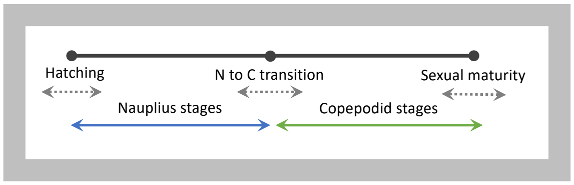


| Individual measurements |
|---|
| ○ Hatching day |
| ○ Copepodid transition day |
| ○ Sexual maturity day |
| ○ Death day |

| Life history stage | Duration (days) |
|---|---|
| egg | ? |
| Nauplius stages | 7-13 |
| ▪ 6 molts | |
| Copepodid stages | 14-29 |
| ▪ 5 molts | |
| Adult | 53-155 |

**B**

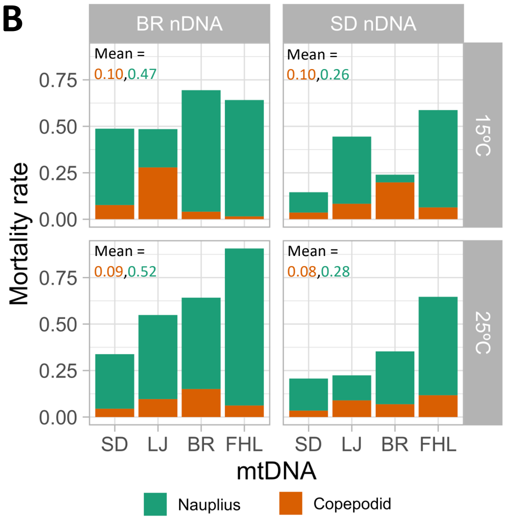


**C**

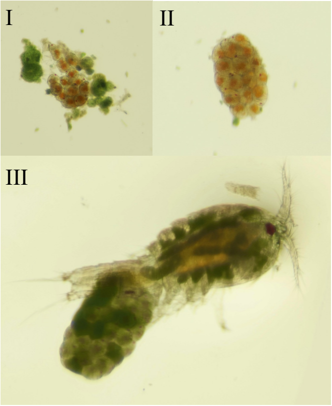


**D**

| Model | Significant terms | | |
|---|---|---|---|
| **Trait: Copepodid development rate** | **Term** | $X^2$ | **P-value** |
| Model: 1 ~ Temp. × nDNA × mtDNA | Intercept | 69.59 | <2.2e-16 |
| + (1\| Family/Clutch) | Temperature | 9.62 | 0.0019 |
| | mtDNA | 11.96 | 0.0176 |
| | mtDNA × nDNA | 11.64 | 0.0203 |
| **Trait: Naupliar development rate** | **Term** | $X^2$ | **P-value** |
| Model: 1 ~ Temp. × nDNA × mtDNA | Intercept | 106.43 | <2.2e-16 |
| + (1\| Family/Clutch) | Temperature | 14.11 | 1.7e-4 |
| | mtDNA × nDNA | 11.62 | 0.0204 |
| **Trait: Total development rate** | **Term** | $X^2$ | **P-value** |
| Model: 1 ~ Temp. × nDNA × mtDNA | Intercept | 142.48 | <2.2e-16 |
| + (1\| Family/Clutch) | Temperature | 19.83 | 8.4e-6 |
| | mtDNA × nDNA | 20.27 | 4.4e-4 |

**Figure S2. Mortality and development rate prior to sexual maturity.** A) Experimental measurements prior to sexual maturity. For each individual, we recorded hatching day, copepodid transition day, sexual maturity day, and death day. From these measurements, we were also able to calculate the duration of the immature stages. Lower values are at 25°C, higher values at 15°C. B) Mortality in immature stages was high, ranging from 14.6% to 90.6% depending on cross and temperature. Naupliar stages had the highest mortality rate, ranging from 4.2% to 84.4%, compared to 1.5% to 27.9% at the copepodid stages. C) Anecdotal evidence across our fertility, longevity, and early mortality experiments suggests that mortality may occur even before hatching (hybrid inviability). In some cases, clutches only partially hatched (I), while in others entire clutches were inviable (II). We also noticed that some clutches were apparently incompletely fertilized, as indicated by the presence of undeveloped green eggs (III). D) Results from type III ANOVA analysis. All models indicate significant influence of temperature and mitonuclear interactions. Naupliar development rate taken as the duration between egg and copepodid transition, while copepodid development rate was taken as the duration between copepodid transition and sexual maturity. Total development rate describes the duration between egg and sexual maturity.
