## Supplementary material for "Mitochondrial effects on fertility and longevity in *Tigriopus californicus* contradict predictions of the mother’s curse hypothesis": Figure S3

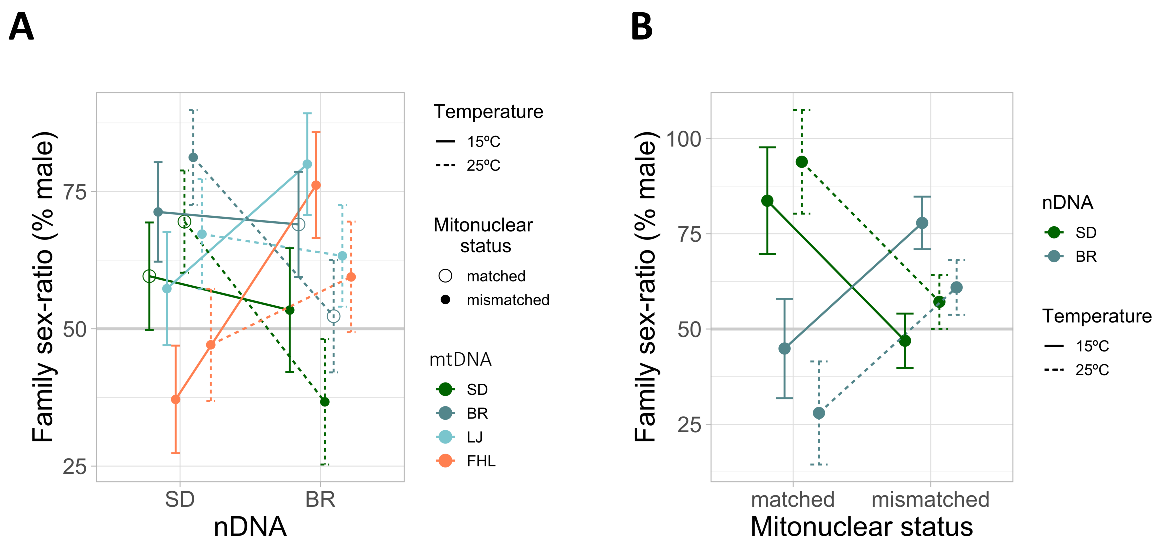


C

| Model | | | Fixed effects | | |
|---|---|---|---|---|---|
| **Model 1. Genotype and environment** | **AICc** | **weight** | **Term** | **$X^2$** | **P-value** |
| Model: ~ Temp. × nDNA + mtDNA × nDNA + (1\| Family) | -42.7 | 0.36 | Intercept | 200.06 | <2.2e-16 |
| | | | Temperature | 22.60 | 2.0e-6 |
| | | | nDNA | 0.11 | 0.7349 |
| | | | mtDNA | 15.89 | 0.0012 |
| | | | Temperature × nDNA | 31.98 | 1.6e-8 |
| | | | mtDNA × nDNA | 29.94 | 1.4e-6 |
| **Model 2. Genotype, environment, and coevolutionary status** | **AICc** | **weight** | **Term** | **$X^2$** | **P-value** |
| Model: ~ mtDNA + Temp. × nDNA + nDNA × status + (1\| Family) | -41.7 | 0.73 | Intercept | 213.00 | <2.2e-16 |
| | | | Temperature | 23.75 | 1.1e-6 |
| | | | nDNA | 13.36 | 2.6e-4 |
| | | | mtDNA | 43.50 | 1.9e-9 |
| | | | Mitonuclear status | 18.51 | 1.7e-5 |
| | | | Temperature × nDNA | 34.41 | 4.5e-9 |
| | | | nDNA x status | 28.77 | 8.1e-8 |

**Figure S3. Family sex ratio modeling.** A) Estimated marginal means (95% CI) from linear mixed-effects model shows significant influence of mitonuclear interactions on mean family sex-ratio. FHL mtDNA has significant effects on family sex ratio depending on nuclear background (mtDNA x nDNA). B) Estimated marginal means (95% CI) from linear mixed-effects model shows significant influence of mitonuclear coevolutionary status on mean family sex ratio. Mismatched status has opposite effects depending on nuclear background (nDNA x status). C) Results from type III ANOVA analysis on family sex ratio models.
