## Supplementary material for "Mitochondrial effects on fertility and longevity in *Tigriopus californicus* contradict predictions of the mother’s curse hypothesis": Document S1

### **Materials and methods**

##### **Inbreeding effects on female hatching number**

In order to measure the impact of male subfertility on female fertility, we conducted a second female fertility experiment in parallel with the first, in which we allowed experimental females to mate with their siblings (as opposed to wild-caught males). Once mated, gravid females were removed to a new dish after they extruded their first egg sac. Gravid females were maintained together until 35 days of age (reared at 25℃) or 70 days of age (reared at 15℃) and then moved into their own dish for fertility assays. Inbreeding female fertility assays were checked MWF until egg sacs hatched, and all offspring were counted under a microscope. For each cross and temperature grouping, we measured the decline in hatching number due to inbreeding by dividing the mean hatching number from inbred crosses from the mean hatching number from outbred crosses (main text).

##### **Mortality and development rate prior to sexual maturity**

In addition to our primary study of adult males and females, we were interested in the impact of mitochondrial variation on mortality and development rate in the naupliar and copepodid stages prior to sexual maturity (Figure S2). To establish families for this analysis, clasped male-female pairs were haphazardly sampled from mitonuclear hybrid or control lines and placed in 60 mm x 15 mm petri dishes filled with ~20 mL of standard culture media, maintained at 20℃ with 12:12 hour light/dark cycle. Dishes were monitored 7 days a week and females were transferred to a fresh dish once gravid with an extruded egg sac. For each experimental family, we alternated moving the first hatched eggsac to either 15℃ or 25℃ so that clutch order was orthogonal to temperature in the experiment. Each family was composed of up to two clutches of fertilized eggs derived from a single mitonuclear hybrid or control female (dam). As clutches hatched, hatching day was recorded and individual nauplii were immediately transferred to 24-well plates containing standard medium. Plates were checked daily for the duration of the experiment, and the copepodid transition day was scored on the day that individual nauplii molted into the first copepodid stage. Nauplii and copepodids are easily distinguished by their size and external morphology. Nauplii have a rounded shape with no segmentation, while individuals in the first copepodid stage have elongated body shape and are almost twice as and faster swimming. Sexual maturity day was scored as in the longevity and fertility experiments. For males, this occurred with the development of geniculate first antennae (claspers) and for females with the development of eggs, which can be seen through the translucent body. Individuals were maintained at the same temperature for the duration of the larval mortality experiment, other than daily inspection under a stereo microscope, which was kept as brief as possible. Death day was recorded as the day that an individual was found unmoving and unresponsive to gentle probing.

##### **Family sex ratio modeling**

Because we recorded the day individuals reached sexual maturity in our longevity experiment, we were able to investigate the influence of genetic and environmental variation on family sex ratio. Family sex ratio was calculated per clutch by counting the number of males and females in our longevity dataset and dividing the number of males from the total number of sexually mature individual in the family. We included a “mortality corrected” estimate of family sex ratio by assigning individuals that died before sexual maturity to the rarer sex in a family. This approach assumes that sex ratio deviations from 50% are due to sex biased mortality, and therefore represents a conservative approach to sex ratio analysis in species with polygenic sex determination. As individuals were not typically transferred to 24-well plates in our longevity dataset until they molted into the copepodid stages, any mortality that occurred before this stage is unknown and uncounted.

We performed linear mixed-effects modeling on family sex ratio and corrected family sex ratio by first using a model selection approach with the *model.sel* function in the MuMIn R package to identify the best fit, most reduced model using an information theoretic approach. Models with weights less than 0.3 were compared using the *anova* function in R in order to identify the best fit model using a chi-square. We included family size as a weighting parameter in all models of family sex ratio and family (replicate) as a random term. Fixed terms included temperature, mtDNA, nDNA as well as temperature x nDNA interactions and mtDNA x nDNA interactions. Models investigating mitonuclear coevolutionary status included temperature, mtDNA, nDNA, mitonuclear coevolutionary status (status) and temperature x nDNA and nDNA x status interactions.

### **Results**

##### **Inbreeding effects on female hatching number**

Inbreeding reduced hatching number in all crosses except for the matched control lines reared at 15℃, which were increased (Figure S1). For the mitonuclear hybrid lines, inbred crosses experienced a 12.5 – 57.65% reduction in hatching number compared to outbred crosses.

The increase in hatching number for inbred vs. outbred matched control crosses could be due to long term laboratory selection, where individuals have been kept at 14-20℃ for several hundred generations. At 25℃, matched control crosses showed a 13.33-32.73% reduction in hatching number due to inbreeding, suggesting that relaxed selection in the laboratory setting may have permitted the accumulation of deleterious mutations that are expressed in higher temperatures.

##### **Mortality and development rate prior to sexual maturity**

As we began our experiment with hatching day, we were not able to record the duration of egg stages. Naupliar stages ranged from 7 days at 25℃ to 13 days at 15℃ on average, and copepodid stages ranged from 14 to 29 days (Figure S2A). From our adult longevity experiment, we found that adult stages ranged from 53-155 days. Mortality rate during the immature stages was very high, ranging from 14.6% to 90.6% mortality depending on cross and temperature (Figure S2B). Naupliar stages had the highest mortality, ranging from 4.2% to 84.4%, while copepodid stages ranged from 1.5% to 27.9% mortality. Mitonuclear hybrid crosses with the FHL mtDNA had among the highest mortality rates, ranging from 58.7% to 90.6% mortality, while the SD control lines had the lowest mortality rates (14.5% at 15℃, 20.7% at 25℃). The latter is a notable contrast with the BR control lines, which had high mortality rates (69.4% at 15℃, and 64.2% at 25℃). Anecdotal evidence across experiments also suggests that mortality may occur even earlier in development. In several experiments, we observed many eggsacs that only partially hatched (I in Figure S2C), with remaining nauplii inviable or apparently unable to emerge from the eggsac. Other observations include several cases where entire fertilized clutches were dropped by the female before hatching (II in Figure S2C), and other cases where eggsacs were incompletely viable (III in Figure S2C). Our linear mixed-effects modeling found significant mtDNA x nDNA interactions for all measured larval traits (copepodid development rate: 𝜒^2^ = 11.64, *p* = 0.0203; naupliar development rate: 𝜒^2^ = 11.62, *p* = 0.0204; and total development rate: 𝜒^2^ = 20.27, *p* = 4.4e-4). Environmental temperature also significantly influenced these traits (copepodid development rate: 𝜒^2^ = 9.62, *p* = 0.0019; naupliar development rate: 𝜒^2^ = 14.11, *p* = 1.7e-4; and total development rate: 𝜒^2^ = 19.83, *p* = 8.4e-6). Increased temperatures resulted in faster developmental rates.

##### **Family sex ratio modeling**

Measures of family sex ratio (percent male) ranged from 0 to 100%, with a mean of 60% and median of 63%. Mean family sex ratio differed among crosses and by temperature, with some mtDNA haplotypes, such as FHL and SD having significantly female biased sex ratios in one nuclear background and male biased in the other (Figure S3A). Mitonuclear status (matched vs. mismatched mitonuclear combinations) also impacted family sex ratio differently depending on nuclear background. Control families with matched nDNA and mtDNA genotypes were male biased for SD nuclear backgrounds and female biased for BR nuclear backgrounds. Introgression of novel “mismatched” mtDNA haplotypes was associated with an increase in the number of males in BR nDNA families and a decrease in the number of males in SD nDNA families (Figure S3B). In BR nDNA lineages, matched families produced an excess of females (compared to 50%), and mismatched mitonuclear hybrids produced an excess of males. SD nDNA control families produced an excess of males and mismatched SD nDNA families produced closer to a 50/50 sex ratio. Temperature effects also impacted the two nuclear lines differently, with SD nDNA families showing more male bias at 25℃ than 15℃, and BR nDNA families showing more female bias at 25℃, which was significant only in mismatched individuals. Linear mixed-effects modeling of family sex ratio (Figure S3C) indicated significant influence of temperature (𝜒^2^ = 22.60, *p* = 2.0e-6), mtDNA haplotype (𝜒^2^ = 15.89, *p* = 0.0012), as well as interactions between mtDNA and nDNA (𝜒^2^ = 29.94, *p* =1.4e-6) and temperature and nDNA (𝜒^2^ = 31.98, *p* = 1.6e-8). Overall, this pattern did not change when using the more conservative mortality-corrected family sex ratio values (Table S4). We also found that family sex ratio was influenced by mitonuclear status (𝜒^2^ = 18.51, *p* = 1.7e-5) as well as nDNA x status interactions (𝜒^2^ = 28.77, *p* = 8.1e-8). Results using mortality-corrected family sex ratio data were consistent with uncorrected family sex ratio (Table S4).

#### **Other observations**

##### **Aberrant clutches in male remating experiments**

In our experiments measuring male remating success, 42 wild type females used in the study produced aberrant clutches when mated to experimental males. This included partially hatched eggs (similar to I in Figure S1C), unhatched fertilized eggs (similar to II in Figure S1C), and partially fertilized eggs (similar to III in Figure S1C). Of these aberrant clutches, 39 were found in crosses between mitonuclear hybrid males and wild-caught females (7.22%) and three were found in control crosses (2.46% of males). Two females, mated with mitonuclear hybrid males, produced aberrant clutches at 15℃, while the remaining 40 females produced aberrant clutches at 25% (95.24% of females producing aberrant clutches). While many wild-caught females died in this experiment, the same proportion of females (16.67%) died in either temperature treatment (15℃: 324/1944; 25℃: 338/2028).
