## Supplementary material for "Mitochondrial effects on fertility and longevity in *Tigriopus californicus* contradict predictions of the mother’s curse hypothesis": Table S1

**Table S1. Sex-effect model selection.** Model selection was performed by ranking models and obtaining model weights. Bolded model terms indicate the term of interest in each model.

| **Trait** | **Temperature** | **Model** | **Rank** | **df** | **AIC_C_** | **ΔAIC** | **Weight** |
| --- | --- | --- | --- | --- | --- | --- | --- |
| Fertility | 15℃ | **Sex x nDNA** + mtDNA | 1 | 9 | 683.6 | 0.00 | 0.934 |
|  |  | **Sex** + mtDNA + nDNA | 2 | 8 | 644.0 | 5.46 | 0.061 |
|  |  | **Sex x mtDNA** + nDNA | 3 | 11 | 649.1 | 10.53 | 0.005 |
|  |  | **Sex x mtDNA x nDNA** | 4 | 18 | 655.9 | 17.35 | 0.000 |
|  | 25℃ | **Sex x mtDNA x nDNA** | 1 | 18 | 1042.0 | 0.00 | 0.991 |
|  |  | **Sex x nDNA** + mtDNA | 2 | 9 | 1052.3 | 10.30 | 0.006 |
|  |  | **Sex** + mtDNA + nDNA | 3 | 8 | 1053.4 | 11.37 | 0.003 |
|  |  | **Sex x mtDNA** + nDNA | 4 | 11 | 1061.6 | 19.64 | 0.000 |
| Longevity | 15℃ | **Sex x mtDNA x nDNA** | 1 | 19 | 15907.2 | 0.00 | 1.000 |
|  |  | **Sex x mtDNA** + nDNA | 2 | 11 | 15945.0 | 37.84 | 0.000 |
|  |  | **Sex x nDNA** + mtDNA | 3 | 9 | 15952.6 | 45.41 | 0.000 |
|  |  | **Sex** + mtDNA + nDNA | 4 | 8 | 15956.1 | 48.93 | 0.000 |
|  | 25℃ | **Sex x mtDNA x nDNA** | 1 | 19 | 12923.6 | 0.00 | 1.000 |
|  |  | **Sex x nDNA** + mtDNA | 2 | 9 | 12959.8 | 36.20 | 0.000 |
|  |  | **Sex x mtDNA** + nDNA | 3 | 11 | 12960.1 | 36.49 | 0.000 |
|  |  | **Sex** + mtDNA + nDNA | 4 | 8 | 12965.7 | 42.05 | 0.000 |
