## Supplementary material for "Mitochondrial effects on fertility and longevity in *Tigriopus californicus* contradict predictions of the mother’s curse hypothesis": Table S2

**Table S2. Sex-effects models.** We performed linear mixed-effects modeling of relative fertility and longevity. Each row represents a separate model, with columns representing modeled fixed effects. We used type III ANOVA to analyze the amount of variance contributing to corrected each trait by the model terms. Cells are marked with a dash when a variable is not included in the model. Values in bold indicate terms with p-values less than α = 0.05.

| **Trait** | **Temperature** | **nDNA** | **mtDNA** | **Sex** | **mtDNAx nDNA** | **Sex x mtDNA** | **Sex x nDNA** | **Sex x mtDNA x nDNA** | **Intercept** |
| --- | --- | --- | --- | --- | --- | --- | --- | --- | --- |
| Fertility | 15℃ | **χ^2^ = 19.92** | **χ^2^ = 9.09** | **χ^2^ = 9.70** | ⎯ | ⎯ | **χ^2^ = 10.09** | ⎯ | **χ^2^ = 160.38** |
|  |  | ***p* = 8.1e-6** | ***p* = 0.0282** | ***p* = 0.0018** |  |  | ***p* = 0.0015** |  | ***p* < 2.2e-16** |
|  | 25℃ | χ^2^ = 0.76 | χ^2^ = 1.34 | χ^2^ = 0.06 | χ^2^ = 0.53 | χ^2^ = 6.09 | χ^2^ = 0.13 | χ^2^ = 9.33 | **χ^2^ = 12.79** |
|  |  | *p* = 0.3848 | *p* = 0.7201 | *p* = 0.8115 | *p* = 0.9131 | *p* = 0.1073 | *p* = 0.7165 | *p* = 0.0252 | ***p* = 3.4e-4** |
| Longevity | 15℃ | χ^2^ = 1.12 | χ^2^ = 6.60 | χ^2^ = 0.54 | χ^2^ = 5.84 | χ^2^ = 2.26 | χ^2^ = 0.70 | χ^2^ = 2.42 | **χ^2^ = 361.18** |
|  |  | *p* = 0.2904 | *p* = 0.0859 | *p* = 0.4609 | *p* = 0.1194 | *p* = 0.5206 | *p* = 0.4033 | *p* = 0.4902 | ***p* < 2.2e-16** |
|  | 25℃ | χ^2^ = 1.47 | **χ^2^ = 14.43** | χ^2^ = 0.86 | **χ^2^ = 10.56** | χ^2^ = 4.94 | χ^2^ = 1.0e-4 | χ^2^ = 5.42 | **χ^2^ = 222.64** |
|  |  | *p* = 0.2247 | ***p* = 0.0024** | *p* = 0.3528 | ***p* = 0.0144** | *p* = 0.1761 | *p* = 0.9912 | *p* = 0.1435 | ***p* < 2.2e-16** |
