## Supplementary material for "Mitochondrial effects on fertility and longevity in *Tigriopus californicus* contradict predictions of the mother’s curse hypothesis": Table S3

**Table S3. Coevolutionary status models**. We performed linear mixed-effects modeling of each trait, comparing a single mtDNA haplotype with both nuclear line controls. The mtDNA column indicates individuals possessing that mtDNA haplotype with both nuclear backgrounds. Each row represents a separate model, with columns representing results from type III ANOVA. All models included the random terms (1 | Family:Clutch) and (1 | nDNA) except for BR and SD mtDNA haplotypes, which do not include the nDNA random term. Cells are marked with a dash when a variable is not included in the model. Shaded cells indicate p-values less than the bonferroni-corrected alpha (95% confidence, p = 0.0167), and bold numbers indicate uncorrected p-values less than 0.05, but greater than the bonferroni-corrected cutoff.

| **Trait** | **mtDNA** | **Temperature** | **Coevolutionary status** | **Intercept** |
| --- | --- | --- | --- | --- |
| Male  remating success | LJ | 𝜒2 = 8.18, p = 0.0042 | 𝜒2 = 0.19, p = 0.6595 | 𝜒2 = 133.89, p < 2.2e-16 |
|  | BR | 𝜒2 =0.17, p = 0.6803 | 𝜒2 = 4.2e-3, p = 0.9481 | 𝜒2 = 61.25, p = 5.0e-15 |
|  | SD | **𝜒2 = 5.40, p = 0.0201** | 𝜒2 = 0.01, p = 0.9150 | 𝜒2 =57.83, p = 2.9e-14 |
|  | FHL | 𝜒2 = 10.55, p = 0.0012 | 𝜒2 = 0.13, p = 0.7151 | 𝜒2 = 49.13, p = 2.4e-12 |
| Female  hatching number | LJ | 𝜒2 = 5.74, p = 0.0165 | 𝜒2 = 0.03, p = 0.8588 | 𝜒2 = 107.80, p < 2.2e-16 |
|  | BR | 𝜒2 = 2.29, p = 0.1305 | 𝜒2 = 1.20, p = 0.2740 | 𝜒2 = 129.85, p < 2.2e-16 |
|  | SD | 𝜒2 = 7.88, p = 0.0050 | 𝜒2 = 6.5e-3, p = 0.9358 | 𝜒2 = 56.09, p = 6.9e-14 |
|  | FHL | 𝜒2 = 6.17, p = 0.0130 | 𝜒2 = 17.90, p = 2.4e-5 | 𝜒2 = 150.25, p < 2.2e-16 |
| Male longevity | LJ | 𝜒2 = 653.21, p < 2.2e-16 | 𝜒2 = 0.09, p = 0.7652 | 𝜒2 = 1064.67, p < 2.2e-16 |
|  | BR | 𝜒2 = 404.84, p < 2.2e-16 | 𝜒2 = 0.09, p = 0.7593 | 𝜒2 = 981.05, p < 2.2e-16 |
|  | SD | 𝜒2 = 266.42, p < 2.2e-16 | 𝜒2 = 9.0e-3, p = 0.9246 | 𝜒2 = 1263.96, p < 2.2e-16 |
|  | FHL | 𝜒2 = 575.13, p < 2.2e-16 | 𝜒2 = 2.54, p = 0.1112 | 𝜒2 = 509.46, p < 2.2e-16 |
| Female longevity | LJ | 𝜒2 = 781.20, p < 2.2e-16 | 𝜒2 = 3.9e-3, p = 0.9500 | 𝜒2 = 2414.79, p < 2.2e-16 |
|  | BR | 𝜒2 = 443.25, p < 2.2e-16 | 𝜒2 = 0.02, p = 0.8836 | 𝜒2 = 1768.77, p < 2.2e-16 |
|  | SD | 𝜒2 = 355.29, p < 2.2e-16 | 𝜒2 = 0.11, p = 0.7443 | 𝜒2 =1085.53, p < 2.2e-16 |
|  | FHL | 𝜒2 = 699.57, p < 2.2e-16 | 𝜒2 = 3.21, p = 0.0733 | 𝜒2 = 346.16, p < 2.2e-16 |
| Naupliar  development rate | LJ | 𝜒2 = 119.43, p < 2.2e-16 | 𝜒2 = 0.76, p = 0.3820 | 𝜒2 = 392.48, p < 2.2e-16 |
|  | BR | 𝜒2 = 72.40, p < 2.2e-16 | 𝜒2 = 1.15, p = 0.2830 | 𝜒2 = 330.31, p < 2.2e-16 |
|  | SD | 𝜒2 = 51.91, p = 5.8e-13 | 𝜒2 = 1.69, p = 0.1934 | 𝜒2 = 254.29, p < 2.2e-16 |
|  | FHL | 𝜒2 = 21.77, p = 3.08e-6 | **𝜒2 =5.72, p = 0.0167** | 𝜒2 = 118.49, p < 2.2e-16 |
| Copepodid development rate | LJ | 𝜒2 = 88.77, p < 2.2e-16 | 𝜒2 = 3.19, p = 0.0740 | 𝜒2 = 273.26, p < 2.2e-16 |
|  | BR | 𝜒2 = 48.32, p = 3.6e-12 | 𝜒2 = 3.1e-3, p = 0.9556 | 𝜒2 = 200.02, p < 2.2e-16 |
|  | SD | 𝜒2 = 59.52, p = 1.2e-14 | 𝜒2 = 1.31, p = 0.2528 | 𝜒2 = 334.12, p < 2.2e-16 |
|  | FHL | 𝜒2 = 42.89, p = 5.8e-11 | **𝜒2 = 5.07, p = 0.0439** | 𝜒2 = 115.87, p < 2.2e-16 |
| Naupliar development rate (Males) | LJ | 𝜒2 = 102.63, p < 2.2e-16 | 𝜒2 = 0.21, p = 0.6464 | 𝜒2 = 498.1, p < 2.2e-16 |
|  | BR | 𝜒2 = 52.45, p = 4.4e-13 | 𝜒2 = 0.72, p = 0.3978 | 𝜒2 = 256.55, p < 2.2e-16 |
|  | SD | 𝜒2 = 71.21, p < 2.2e-16 | 𝜒2 = 0.29, p = 0.5889 | 𝜒2 = 278.99, p < 2.2e-16 |
|  | FHL | 𝜒2 = 53.71, p = 2.3e-13 | 𝜒2 = 1.16, p = 0.2820 | 𝜒2 = 288.56, p < 2.2e-16 |
| Naupliar development rate (Females) | LJ | 𝜒2 = 68.84, p < 2.2e-16 | 𝜒2 =0.65, p = 0.4204 | 𝜒2 = 282.91, p < 2.2e-16 |
|  | BR | 𝜒2 = 87.82, p < 2.2e-16 | 𝜒2 = 1.57, p = 0.2102 | 𝜒2 = 387.47, p < 2.2e-16 |
|  | SD | 𝜒2 = 19.59, p = 9.6e-6 | 𝜒2 = 1.67, p = 0.1967 | 𝜒2 = 81.98, p < 2.2e-16 |
|  | FHL | 𝜒2 = 13.43, p = 0.0002 | **𝜒2 =5.20, p = 0.0225** | 𝜒2 = 66.66, p = 3.2e-16 |
| Copepodid development rate (Males) | LJ | 𝜒2 =128.11, p < 2.2e-16 | 𝜒2 =1.43, p = 0.2319 | 𝜒2 =369.59, p < 2.2e-16 |
|  | BR | 𝜒2 =50.68, p = 1.1e-12 | 𝜒2 =4.7e-3, p = 0.9451 | 𝜒2 =199.60, p < 2.2e-16 |
|  | SD | 𝜒2 =94.61, p < 2.2e-16 | 𝜒2 =1.02, p = 0.3117 | 𝜒2 =552.98, p < 2.2e-16 |
|  | FHL | 𝜒2 =33.04, p = 9.0e-9 | 𝜒2 =1.52, p = 0.2182 | 𝜒2 =202.52, p < 2.2e-16 |
| Copepodid development rate (Females) | LJ | 𝜒2 =42.78, p = 6.1e-11 | 𝜒2 =3.05, p = 0.0806 | 𝜒2 =112.45, p < 2.2e-16 |
|  | BR | 𝜒2 =24.54, p = 7.3e-7 | 𝜒2 =0.13, p = 0.7152 | 𝜒2 =100.48, p = 7.3e-7 |
|  | SD | 𝜒2 =20.05, p = 7.5e-6 | 𝜒2 =0.08, p = 0.7836 | 𝜒2 =104.51, p < 2.2e-16 |
|  | FHL | 𝜒2 =41.54, p = 1.2e-10 | **𝜒2 =4.39, p = 0.0362** | 𝜒2 =150.88, p < 2.2e-16 |
| * Models required a bound optimization by quadratic approximation (bobyqa), allowing up to 2e5 iterations in order to converge. | | | | |
