## Supplementary material for "Mitochondrial effects on fertility and longevity in *Tigriopus californicus* contradict predictions of the mother’s curse hypothesis": Table S4

**Table S4. Mortality-corrected family sex ratio.** We performed linear mixed-effects modeling on mortality-corrected family sex ratio using two formulas. We used type III ANOVA to analyze the amount of variance contributing to corrected family sex ratio by the model terms.

| **Model 1: Genotype and environment** | **Model Term** | **χ^2^** | **P-value** |
| --- | --- | --- | --- |
| ~ Temperature x nDNA + mtDNA x nDNA + | Intercept | 219.74 | <2.2e-16 |
| (1\| Family) | Temperature | 16.58 | 4.7e-5 |
|  | nDNA | 0.19 | 0.6667 |
|  | mtDNA | 11.03 | 0.0116 |
|  | Temperature x nDNA | 31.00 | 2.6e-8 |
|  | mtDNA x nDNA | 14.53 | 0.0023 |
| **Model 2: Genotype, environment, and coevolutionary status** | **Model Term** | **χ^2^** | **P-value** |
| ~ mtDNA + Temperature x nDNA + nDNA x | Intercept | 237.65 | <2.2e-16 |
| coevolutionary status + (1\| Family) | Temperature | 16.43 | 5.0e-5 |
|  | nDNA | 5.20 | 0.0226 |
|  | mtDNA | 17.77 | 4.9e-4 |
|  | Coevolutionary status | 10.78 | 0.0010 |
|  | Temperature x nDNA | 30.54 | 3.3e-8 |
|  | nDNA x status | 12.67 | 3.7e-4 |
