## Supplementary material for "Mitochondrial effects on fertility and longevity in *Tigriopus californicus* contradict predictions of the mother’s curse hypothesis": Table S5

**Table S5. Best-fit models.** We performed linear mixed-effects modeling of sex-specific fertility and longevity. Each row represents a separate model, with columns representing modeled fixed effects. We used type III ANOVA to analyze the amount of variance contributing to each trait by the model terms. Cells are marked with a dash when a variable is not included in the model. Values in bold indicate terms with p-values less than α = 0.05. Abbreviations: Temp. = Temperature

| **Trait** | **Temperature** | **nDNA** | **mtDNA** | **mtDNA x nDNA** | **Temp. x nDNA** | **Temp. x mtDNA** | **Temp. x mtDNA x nDNA** | **Intercept** |
| --- | --- | --- | --- | --- | --- | --- | --- | --- |
| Male remating success | **χ^2^ = 5.05** | χ^2^ = 0.15 | χ^2^ = 0.29 | χ^2^ = 2.33 | **χ^2^ = 7.41** | χ^2^ = 6.37 | **χ^2^ = 16.95** | **χ^2^ = 53.73** |
|  | ***p* = 0.0246** | *p* = 0.6991 | *p* = 0.9621 | *p* = 0.5053 | ***p* = 0.0065** | *p* = 0.0949 | ***p* = 7.2e-4** | ***p* <2.3e-13** |
| Female hatching number | **χ^2^ = 12.42** | χ^2^ = 1.37 | **χ^2^ = 20.36** | ⎯ | ⎯ | ⎯ | ⎯ | **χ^2^ = 110.10** |
|  | ***p* = 4.2e-4** | *p* = 0.2416 | ***p* = 1.4e-4** |  |  |  |  | ***p* <2.2e-16** |
| Male longevity | **χ^2^ = 681.28** | **χ^2^ = 20.36** | χ^2^ = 5.57 | ⎯ | **χ^2^ = 4.19** | ⎯ | ⎯ | **χ^2^ = 2317.0** |
|  | ***p* <2.2e-16** | ***p* = 6.4e-6** | *p* = 0.1348 |  | ***p* = 0.0407** |  |  | ***p* <2.2e-16** |
| Female longevity | **χ^2^ = 638.01** | **χ^2^ = 8.31** | **χ^2^ = 14.66** | **χ^2^ = 12.03** | **χ^2^ = 4.05** | ⎯ | ⎯ | **χ^2^ = 1149.1** |
|  | ***p* < 2.2e-16** | ***p* = 0.0040** | ***p* = 0.0021** | ***p* = 0.0073** | ***p* = 0.0442** |  |  | ***p* <2.2e-16** |
