## Supplementary material for "Mitochondrial effects on fertility and longevity in *Tigriopus californicus* contradict predictions of the mother’s curse hypothesis": Table S6

**Table S6. Survivorship analysis post hoc significance tests.** We performed parametric survivorship regression, using the Gompertz model to identify the impacts of temperature, nDNA, mtDNA, sex, and two-way interactions on frailty (lnα) and rate of senescence (β). We used Kruskal-Wallis one-way analysis of variance to estimate the significance of parameter estimates. Values in bold indicate terms with p-values less than α= 0.05. Cells are marked with a dash when a variable is not included in the model. Abbreviations: Temp. = Temperature

| **Parameter** | **Temp.** | **nDNA** | **mtDNA** | **Sex** | **Temp. x mtDNA** | **Temp. x nDNA** | **Temp. x Sex** | **mtDNA x nDNA** | **nDNA x Sex** |
| --- | --- | --- | --- | --- | --- | --- | --- | --- | --- |
| Frailty | **χ^2^ = 22.55** | χ^2^ = 0.89 | χ^2^ = 0.53 | χ^2^ = 1.11 | **χ^2^ = 23.09** | **χ^2^ = 23.76** | **χ^2^ = 24.83** | χ^2^ = 4.10 | χ^2^ = 2.24 |
|  | ***p* = 2.04e-6** | *p* = 0.3461 | *p* = 0.9132 | *p* = 0.2913 | ***p* = 0.0016** | ***p* = 2.81e-5** | ***p* = 1.98e-5** | *p* = 0.7679 | *p* = 0.5238 |
| Frailty | 15℃ | **χ^2^ = 3.98** | χ^2^ = 0.82 | χ^2^ = 0.04 | ⎯ | ⎯ | ⎯ | χ^2^ = 12.53 | χ^2^ = 4.13 |
|  |  | ***p* = 0.0460** | *p* = 0.8456 | *p* = 0.8336 |  |  |  | *p* = 0.0844 | *p* = 0.2483 |
| Frailty | 25℃ | χ^2^ = 0.40 | χ^2^ = 1.15 | **χ^2^ = 6.89** | ⎯ | ⎯ | ⎯ | χ^2^ = 4.06 | **χ^2^ = 8.18** |
|  |  | *p* = 0.5286 | *p* = 0.7657 | ***p* = 0.0074** |  |  |  | *p* = 0.7730 | ***p* = 0.0424** |
| Frailty | 15℃ | BR | χ^2^ = 5.17 | χ^2^ = 0.08 | ⎯ | ⎯ | ⎯ | ⎯ | ⎯ |
|  |  |  | *p* = 0.1600 | *p* = 0.7728 |  |  |  |  |  |
| Frailty | 15℃ | SD | χ^2^ = 6.00 | χ^2^ = 0.00 | ⎯ | ⎯ | ⎯ | ⎯ | ⎯ |
|  |  |  | *p* = 0.1116 | *p* = 1.00 |  |  |  |  |  |
| Frailty | 25℃ | BR | χ^2^ = 2.67 | χ^2^ = 1.33 | ⎯ | ⎯ | ⎯ | ⎯ | ⎯ |
|  |  |  | *p* = 0.4459 | *p* = 0.2482 |  |  |  |  |  |
| Frailty | 25℃ | SD | χ^2^ = 1.17 | **χ^2^ = 5.33** | ⎯ | ⎯ | ⎯ | ⎯ | ⎯ |
|  |  |  | *p* = 0.7610 | ***p* = 0.0209** |  |  |  |  |  |
| Senescence | **χ^2^ = 23.27** | χ^2^ = 2.88 | χ^2^ = 0.41 | χ^2^ = 1.28 | **χ^2^ = 23.78** | **χ^2^ = 26.26** | **χ^2^ = 24.83** | χ^2^ = 4.46 | χ^2^ = 4.33 |
|  | ***p* = 1.41e-6** | *p* = 0.0899 | *p* = 0.9384 | *p* = 0.2582 | ***p* = 0.0012** | ***p* = 8.40e-6** | ***p* = 1.68e-5** | *p* = 0.7255 | *p* = 0.2283 |
| Senescence | 15℃ | **χ^2^ = 8.04** | χ^2^ = 0.73 | χ^2^ = 0.71 | ⎯ | ⎯ | ⎯ | χ^2^ = 12.22 | **χ^2^ = 8.85** |
|  |  | ***p* = 0.0046** | *p* = 0.8666 | *p* = 0.4008 |  |  |  | *p* = 0.0935 | ***p* = 0.0314** |
| Senescence | 25℃ | χ^2^ = 3.57 | χ^2^ = 1.26 | **χ^2^ = 5.34** | ⎯ | ⎯ | ⎯ | χ^2^ = 6.75 | **χ^2^ = 9.62** |
|  |  | *p* = 0.0587 | *p* = 0.7393 | ***p* = 0.0209** |  |  |  | *p* = 0.4454 | ***p* = 0.0221** |
| Senescence | 15℃ | BR | χ^2^ = 3.50 | χ^2^ = 0.33 | ⎯ | ⎯ | ⎯ | ⎯ | ⎯ |
|  |  |  | *p* = 0.3208 | *p* = 0.5637 |  |  |  |  |  |
| Senescence | 15℃ | SD | χ^2^ = 4.83 | χ^2^ = 0.75 | ⎯ | ⎯ | ⎯ | ⎯ | ⎯ |
|  |  |  | *p* = 0.1844 | *p* = 0.3865 |  |  |  |  |  |
| Senescence | 25℃ | BR | χ^2^ = 1.00 | χ^2^ = 3.00 | ⎯ | ⎯ | ⎯ | ⎯ | ⎯ |
|  |  |  | *p* = 0.8013 | *p* = 0.0833 |  |  |  |  |  |
| Senescence | 25℃ | SD | χ^2^ = 1.67 | **χ^2^ = 5.33** | ⎯ | ⎯ | ⎯ | ⎯ | ⎯ |
|  |  |  | *p* = 0.6444 | ***p* = 0.0209** |  |  |  |  |  |
