## Supplementary material for "Mitochondrial effects on fertility and longevity in *Tigriopus californicus* contradict predictions of the mother’s curse hypothesis": Table S7

**Table S7. Contribution of mitochondrial genetic variance to fertility and longevity.** For mitochondrial coefficient of variation (CV_mt_), we present the median difference between sexes, the 95% confidence interval, p-value, and indicate which sex exhibited higher mitochondrial genetic variance for each grouping.

| **Trait** | **Temperature** | **nDNA** | **CV_mt_** |
| --- | --- | --- | --- |
| Fertility | 15℃ | BR | 0.1061 |
|  |  |  | 0.0971, 0.1131 |
|  |  |  | *p* < 2.2e-16 |
|  |  |  | Females > Males |
|  |  | SD | 0.1032 |
|  |  |  | 0.0967, 0.1106 |
|  |  |  | *p* < 2.2e-16 |
|  |  |  | Females > Males |
|  | 25℃ | BR | -0.0993 |
|  |  |  | -0.1078, -0.0930 |
|  |  |  | *p* < 2.2e-16 |
|  |  |  | Males > Females |
|  |  | SD | 0.0076 |
|  |  |  | 0.0007, 0.0147 |
|  |  |  | *p* = 0.0209 |
|  |  |  | Females > Males |
| Longevity | 15℃ | BR | 0.0230 |
|  |  |  | 0.0202, 0.0250 |
|  |  |  | *p* < 2.2e-16 |
|  |  |  | Females > Males |
|  |  | SD | 0.0039 |
|  |  |  | 0.0015, 0.0062 |
|  |  |  | *p* < 2.2e-16 |
|  |  |  | Females > Males |
|  | 25℃ | BR | 0.0313 |
|  |  |  | 0.0291, 0.0335 |
|  |  |  | *p* < 2.2e-16 |
|  |  |  | Females > Males |
|  |  | SD | 0.0488 |
|  |  |  | 0.0463, 0.0522 |
|  |  |  | *p* < 2.2e-16 |
|  |  |  | Females > Males |
